# Improved estimates of abortion rates in tsetse (*Glossina* spp)

**DOI:** 10.1101/2022.09.15.508167

**Authors:** J. W. Hargrove

## Abstract

Abortion rates were assessed for 170, 846 tsetse (154,228 *G. pallidipes* and 19,618 *G. m. morsitans*) sampled in Zimbabwe in 1988 - 1999. Abortions were diagnosed for flies where the uterus was empty and the largest oocyte awaiting ovulation was <82% of the expected mature length. Of tsetse caught in odour-baited traps, 0.64% (95% ci: 0.59 - 0.69) of *G. pallidipes* and 1.00% (0.76-1.29) of *G. m. morsitans* were diagnosed as having suffered a recent abortion. For flies from artificial refuges, abortion rates were higher, 2.03% (1.77 - 2.31) and 2.28% (1.85 - 2.79) for the two species, respectively. Abortion rates decreased with increasing wing fray and contrary to laboratory findings, did not increase in the oldest flies; they were highest in the hottest months and years, and increased with decreasing adult wing length. Percentages of tsetse with empty uteri, regardless of abortion status, were significantly higher than the estimated abortion percentages. For tsetse from traps, 4.01% (95% ci: 3.90 – 4.13) of *G. pallidipes* and 2.52% (2.14-2.95) of *G. m. morsitans* had empty uteri; for flies from artificial refuges the percentage were 12.69% (12.07 – 13.34) and 14.90% (13.82 -16.02), respectively.

## Introduction

For tsetse (*Glossina* spp), vectors of the African animal and human trypanosomiases, each birth event consists of the deposition of a single larva of similar weight to its mother (English *et al*., 2016; Hargrove *et al*., 2018; Haines *et al*., 2020). Unsurprisingly, larviposition only occurs at intervals of about nine days, depending on temperature. Given this low birth rate, tsetse populations can only persist if mortality rates are correspondingly small (Vale, 1974; Kajunguri *et al*., 2019). Losses of order 95% as reported for mosquitoes (Mogi *et al*., 1984) would not be sustainable in a tsetse population (Hargrove, 1988). The deposited larva does not feed, but burrows into the ground where it is deposited and forms around itself a hard puparial shell, inside which all further pre-adult development. These stages are thus better protected than immature mosquitoes, both from the elements and from biological enemies, although they are still subject to parasitism (Heaversedge, 1969) and predation (Rogers & Randolph, 1984).

The egg and larval instars are likewise protected in the maternal uterus. They are thus as safe as the mother that feeds them, but if the mother fails to ingest sufficient blood, her own health and that of her larva will be jeopardised and she may abort the developing offspring. Madubunyi (1975, 1978) suggested a method for identifying abortions among tsetse, known to have ovulated at least once, where the uterus is empty. Such flies have either; (i) recently larviposited and not yet ovulated again, or (ii) aborted either an egg or early-stage larva. In the laboratory, about 50% of laboratory *Glossina m. morsitans* Westwood ovulated within 1 hour of larviposition, and 80% within two hours (Denlinger & Ma, 1974). The expected length (*la*) of the largest oocyte in the ovaries of a normal postpartum should thus be approximately the same length (*le*) as the egg – whose size does not increase while it is *in utero*. If, however, the fly has aborted in early pregnancy, we expect *la* << *le* and Madubunyi (1975) identified an abortion where a female had an empty uterus and where *la* < *le* - 1.96 × standard deviation (*le*).

Turner & Snow, (1984), Okiwelu (1977), Van Sickle & Phelps (1988) and Hargrove (1999a) have used this approach with which, however, there at two problems. If we dissect a female shortly after she aborted early in pregnancy, we expect the uterus to be empty and the largest oocyte to be much smaller than its mature size. We will then, correctly, diagnose an abortion. If, however, the same fly were captured five or six days after the abortion, the fly might still not have ovulated again – but the oocyte would now be much closer to its mature size, and we might then, incorrectly, diagnose the fly as postpartum. The other problem is that, in the studies cited above, there was no way of checking whether the diagnosis of an abortion, or lack of one, was correct.

The first problem could result in under-estimates of abortion rates. Hargrove (1999a) estimated abortion rates of order 1% among 140,000 female *G. m. morsitans* and *G. pallidipes* Austen, dissected in Zimbabwe between 1988 and 1995. He noted, however, that the frequencies of empty uteri in dissected flies was significantly higher than the estimated frequency of abortions. A further 34,000 tsetse were dissected between 1995 and 1999 and the augmented data set is analysed here to reassess the questions raised above.

Included in the new set are dissection results for females captured, in artificial warthog burrows, just prior to larviposition (Muzari & Hargrove, 2005; Hargrove *et al*., 2018). This allowed the capture of full-term-pregnant tsetse, and measurements on oocytes of flies with empty uteri, which had just produced a larva/pupa, whose linear dimensions and weight could also be determined. This allowed measurement, in manifestly postpartum females, of egg lengths in females that had ovulated prior to dissection, or the largest oocyte where they had not yet ovulated. The aim is to use the larger sample, and new information from the burrows, to produce improved estimates of abortion rates and to elucidate the factors affecting these rates – particularly the effects of fly age and size and of weather, particularly temperatures, that females experience during pregnancy.

## Methods

### Fly sampling

Tsetse were captured at Rekomitjie Research Station, Zambezi Valley, Zimbabwe, 16° 10′S 29° 25′ E, altitude 503 m, between September 1988 and December 1999, using; (i) odour-baited mechanical traps (Hargrove & Langley, 1990; (ii) artificial refuges (Vale, 1971); (iii) electrical devices (Vale, 1974); (v) hand nets, used to catch flies from stationary and mobile oxen (Pilson & Leggate, 1962); (v) artificial warthog burrows (Muzari & Hargrove, 1995). Hargrove *et al*. (2019) describe these devices and their use, and the ovarian dissection technique. We follow Randolph *et al*. (1991) in assigning larvae to the third instar only if they had blackened polypneustic lobes, *i*.*e*., those that are in approximately the last day of pregnancy (Denlinger & Ma, 1974). Larvae classified as second instar thus also include larvae in the early and middle part of their third instar.

### Estimating the expected length of a mature oocyte

The multiple linear regression approach of Hargrove (1999a) was extended to include further independent variables affecting the expected egg length (*elhat*) of an egg produced by a female tsetse. These included her ovarian age and wing length, and the mean temperature over the nine days prior to her capture. For a normal postpartum fly, the length (*al*) of the largest oocyte in such flies should be close to *elhat*. The results of Hargrove *et al*. (2018) were used to estimate a cut-off value of the quotient *q* = *al*/*elhat* to separate, among females with empty uteri, those flies that had recently deposited a full-term larva and those that had experienced an abortion. In that study, full-term pregnant tsetse were captured when they entered artificial warthog burrows prior to larviposition. The burrows were cleared at approximately 90-minute intervals and most females were found with a late third instar *in utero*. Smaller proportions deposited a full-term late third instar larva in the burrow: on dissection these flies were generally found to have an empty uterus but, by definition, had not experienced an abortion. Accordingly, the above-mentioned cut-off value of *q* was set at the level below which no fly with an empty uterus was found to have produced a pupa.

### Detecting abortions

Abortion rates were measured only among those flies where a dissection was completed to the point where the ovarian category could be ascertained. Flies in ovarian category 0 had never been pregnant and were excluded from the analysis. For older flies, abortion could only be assessed if the dissector reported that the uterus was indeed empty, and also provided a measure for the largest oocyte. Failure to satisfy these criteria occurred in only 0.1% of the flies eligible for abortion analysis; 23/17,788 of *G. m. morsitans* and 162/142,081 of *G. pallidipes*.

### Measurements of meteorological data

Daily maximum and minimum temperatures were recorded from a mercury thermometer in a Stevenson screen at Rekomitjie Research Station. A rain gauge, placed about 3 metres from the screen, was used to measure daily rainfall.

### Statistical analyses

Data, and statistical, analyses were carried out using Microsoft Excel and StataCorp Stata, Version 14.2. All error limits in this report indicate 95% confidence intervals. Stepwise multiple linear regression was used to identify the factors that determine the expected egg length (*elhat*) in *G. m. morsitans* and *G. pallidipes*. Logistic regression was used to identify the factors likely associated with abortion: ovarian and wing fray categories, month and year of capture, and ambient temperature around the time that a fly was captured. When using year of capture as a factor, years were taken to start on 1 July of a given calendar year, and to end on 30 June of the following calendar year. This ensured that each rainy season always fell entirely in a year of measure. Factors were added singly in a stepwise fashion, with each additional factor only being added to the model if application of a likelihood ratio (LR) test indicated a significant effect at the 0.05 level of probability. Results of logistic regressions were used to predict abortion probabilities using Stata’s “*margin*” command. These statistics were calculated at fixed values of some covariates and specified – or averaged – values of the other covariates in the model.

### Properties of female tsetse processed

Table S1.1 in the Supplementary File S1 provides details for all flies collected and processed at Rekomitjie between September 1988 and December 1999. Males and pupae were excluded from the present analyses, as were flies subjected only to nutritional analysis, or where dissection was carried out only to assess whether the fly was infected with trypanosomes. For 5.6% of the remaining *G. m. morsitans*, and 1.9% of the *G. pallidipes*, flies were damaged during collection and could not be dissected. Most damaged flies, 77% of *G. m. morsitans* and 55% of *G. pallidipes*, were collected using electric nets. Flies sticking to the electric grid were dried out or were killed and sometimes decomposed before they could be dissected. Flies captured in mechanical traps were not subject to these problems. A much smaller proportion of the flies processed, 1.0% of *G. m. morsitans* and 0.5% for *G. pallidipes*, were damaged during dissection, to the point that the ovarian category of the fly could not be ascertained. All damaged flies were excluded from analyses of abortion rates, as were 86/19,618 (0.4%) of all *G. m. morsitans* and 326/154,228 of *G. pallidipes* (0.2%) where a major reproductive anomaly was diagnosed (Turner & Snow, 1984; Hargrove, 1999a). Flies more than 9 days old but still not inseminated (29/37,862 = 0.05% for *G. pallidipes* and 2/3787 = 0.05% for *G. m. morsitans*) were excluded from further analysis. For *G. pallidipes*, 25/29 of these flies were recorded in the hot months of September-February.

## Results

### Expected egg length

The full results of multiple regression modelling of egg length against various independent variables are shown in Supplementary File S1, Tables S1.2A and S1.2B for *G. pallidipes* and *G. m. morsitans*, respectively. Expected egg length increased with maternal wing length and decreased with mean daily maximum temperature over the nine days prior to capture: both were used as continuous variables, and both had statistically significant effects at the 0.01 level of probability (*t* test). Expected egg length varied significantly with month and year of capture, and maternal ovarian category – entered into the regression as categorical variables (Figure 1). For both species, egg length was shortest in November at the end of the hot dry season, and longest in April-May, after the end of the rainy season (Fig. 1A, C). For *G. pallidipes*, egg length declined between 1988 and 1992, and increased thereafter: the pattern was not clearly seen in *G. m. morsitans* – perhaps due in part to the smaller sample sizes (Fig. 1B, E). For both species, egg lengths were markedly shorter in mothers in their first pregnancy. For *G. pallidipes*, eggs were also significantly shorter in ovarian category 2 than for older flies (Figure 1C): for *G. m. morsitans* the same trend was seen, although the egg length for flies in ovarian category 2 was not significantly shorter than in older flies (Fig. 1F).

**Figure 1.**
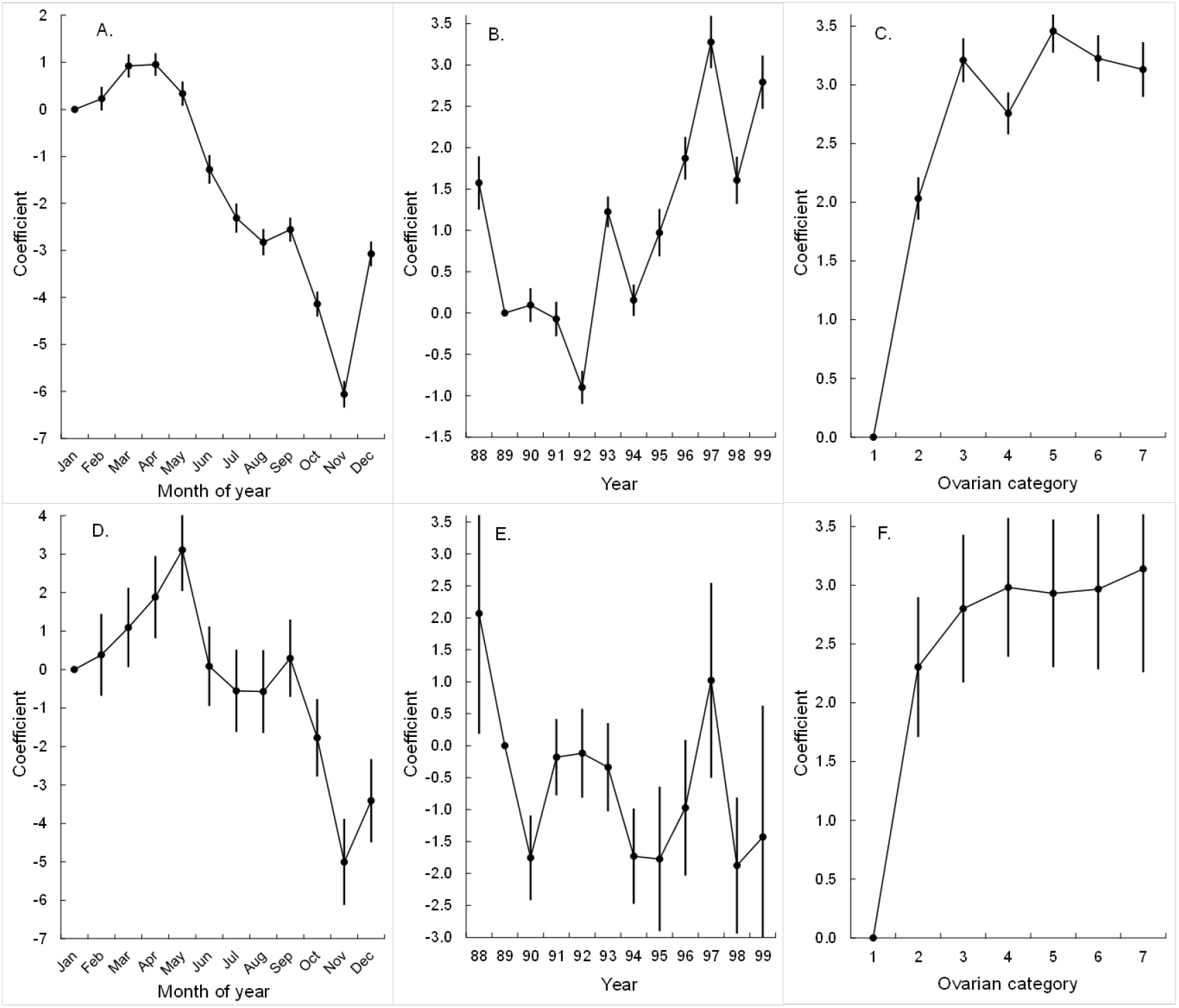
Values of coefficients estimated by multiple linear regression of observed egg lengths in *G. pallidipes* (A, B, C) and *G. m. morsitans* (D, E, F) as functions of: A. Capture month (baseline January); B. Capture year (baseline 1989); C. ovarian category of the mother (baseline ovarian category = 1). Sample sizes, *N* = 67,057 and 6244, respectively. Egg length also increased with increasing wing length and decreased with increasing maximum temperature over the nine days prior to the capture of the fly. Full results of regression analyses in Supplementary File S1, Tables S1.2A and S1.2B.

### Oocyte length as a fraction of expected egg length

#### Pregnant flies

When there is an egg *in utero*, the largest oocyte in the ovaries is about 35% of the expected egg length; this value increases to about 75% during the time when there is a first instar larva *in utero* (Fig. 2). The modal oocyte length among flies with a second instar larvae *in utero* is already of the same order as the expected egg length, and there is little change in oocyte length for flies with a third instar larva *in utero*. These results provide a rough guide to the presumed uterine content aborted by tsetse found to have empty uteri.

**Figure 2.**
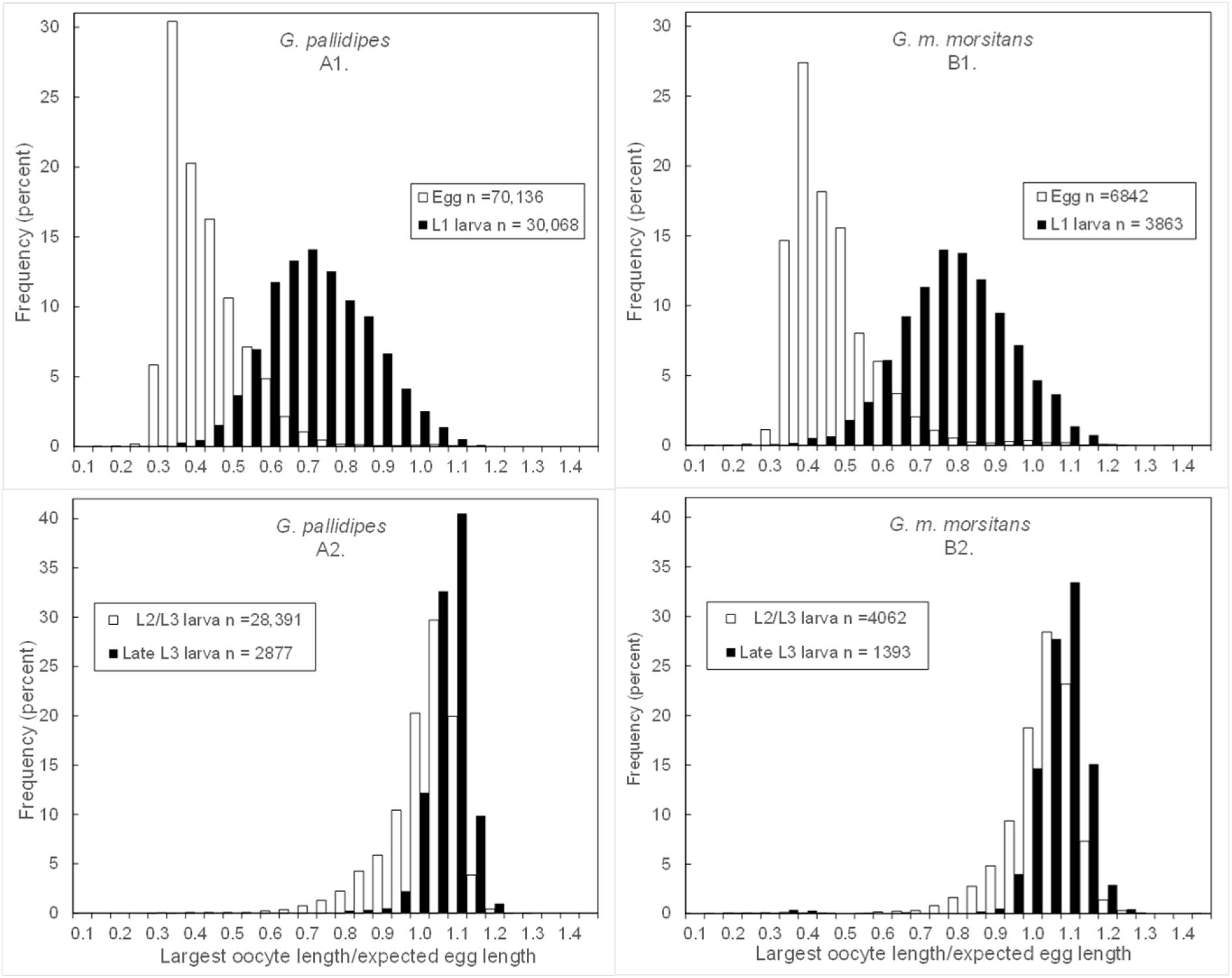
Oocyte lengths, as a quotient of expected egg length, in female *G. m. morsitans* and *G. pallidipes* with different uterine inclusions.

### Flies with empty uteri

Expected egg lengths, *elhat*, for both species show a unimodal distribution skewed towards smaller flies (Fig 3A, B). For flies with empty uteri, *q* = *al*/*elhat* has a major peak with a modal value just less than 1.0 (Fig 3C, D). Thus, as expected for flies that have completed a normal pregnancy, the size of the largest oocyte is close to the size of the egg that is about to be ovulated. To the left of this peak there is an approximately uniform distribution of flies with values of the quotient *q* = *al*/*elhat* ranging from 0.2-0.8, consistent with a large proportion of these flies having recently aborted an egg or a first instar larva (*cf* Figure 2).

**Figure 3.**
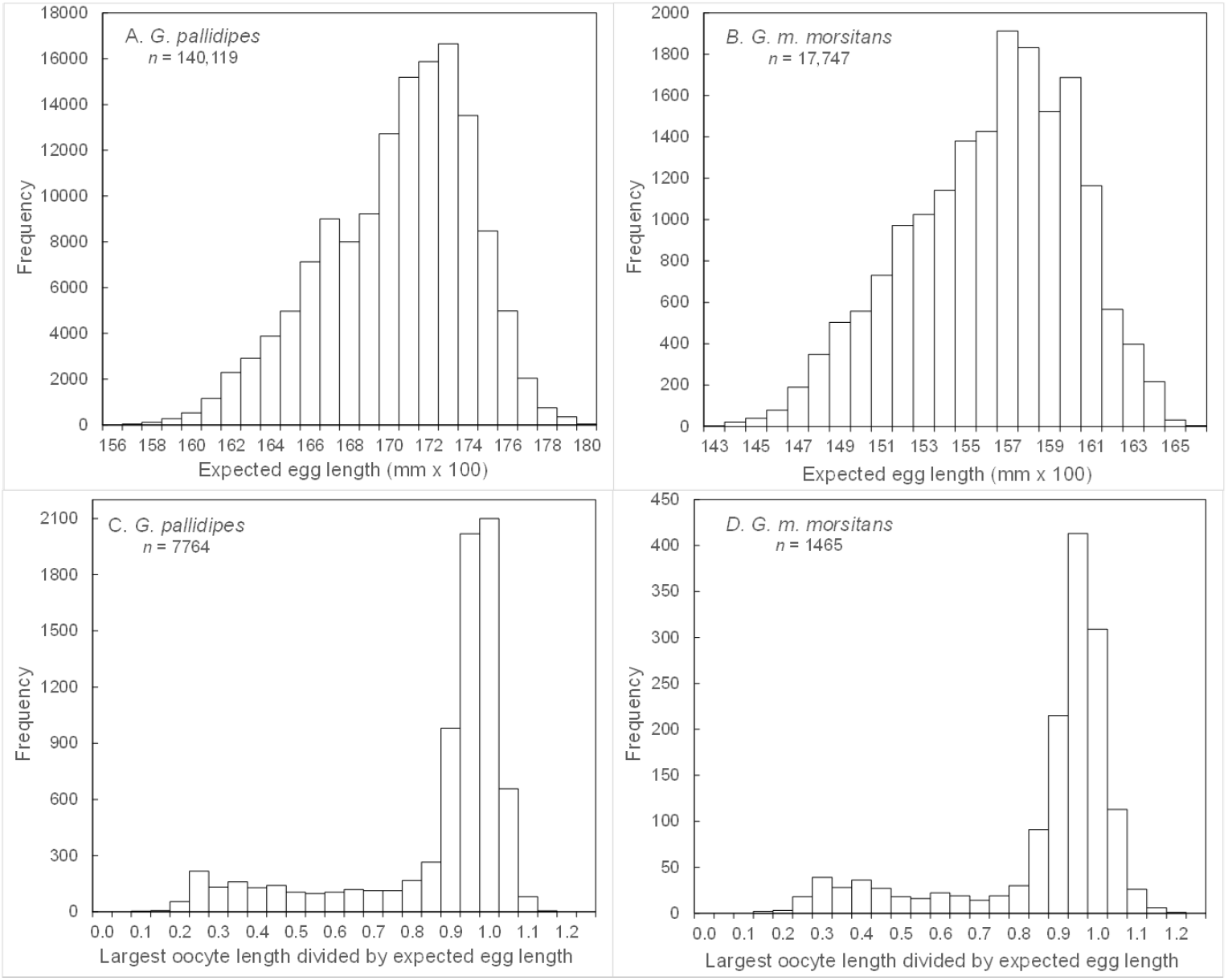
Distributions of expected egg lengths (*elhat*) for A. *G. pallidipes* and B. *G. m. morsitans* – as calculated from the regression results (Table S1.2A and 2B). Distribution of the quotient *q* = *al*/*elhat* for C. *G. pallidipes* and D. *G. m. morsitans*, calculated only for flies that had ovulated at least once and where the uterus was empty. Notice the differences in scales.

### Criteria for diagnosing an abortion

#### i) Empty uterus

The results of Denlinger & Ma (1974) suggest that, for *G. m. morsitans* in the laboratory, about 80% of females ovulate within two hours of depositing a larva. The burrows data (Hargrove *et al*., 2018) provides a different picture for field-collected flies; of 27 *G. pallidipes* that produced a fully mature larva, and were dissected on the following day, none had yet ovulated. Moreover, of postpartum flies dissected 2-6 hours after capture, only 34/526 = 6.5% of *G. pallidipes*, and 2/175 (1.1%) of *G. m. morsitans*, had ovulated.

#### ii)Largest oocyte in ovaries smaller than expected

An appropriate cut-off value of *q*, for diagnosing an abortion, must clearly lie somewhere to the left of the major peaks in Figures 3C for *G. pallidipes* and 3D for *G. m. morsitans*. Figure 4A shows the proportions of female *G. pallidipes* seen to have produced a pupa, among those sampled in artificial warthog burrows, which were found to have an empty uterus. For values of *q* ≥ 0.82, more than 40% of flies captured in artificial warthog burrows were observed to have produce a full-term viable larva and, by definition, had not suffered a recent abortion.

**Figure 4.**
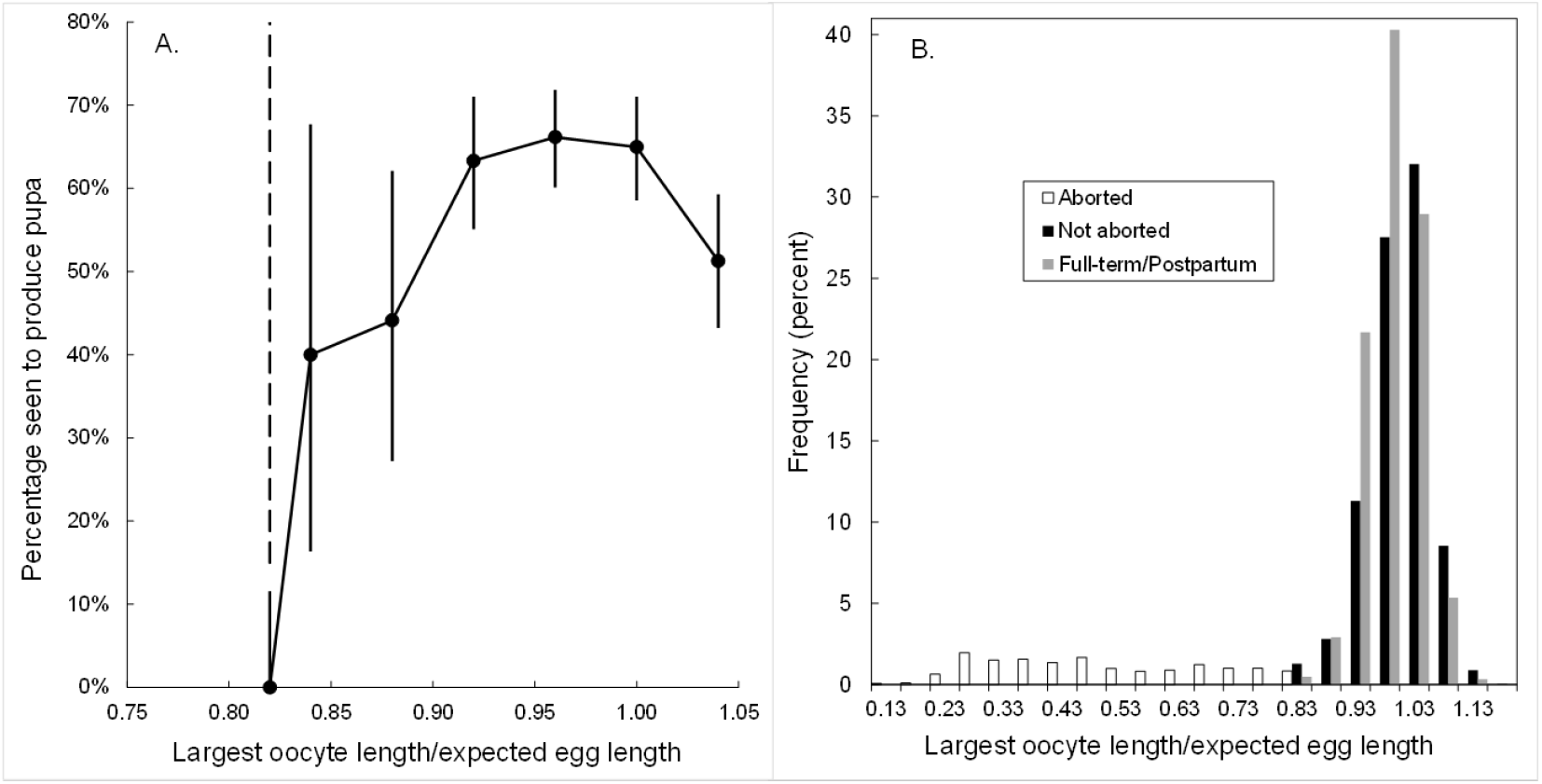
A. Proportions of female *G. pallidipes* seen to have produced a pupa, among those sampled in artificial warthog burrows, which were found to have an empty uterus (*n* = 4057). Plotted as a function of the quotient *q* = *al*/*elhat*. B. Distribution of the quotient *q* = *al*/*elhat* for *G. pallidipes* caught in traps among those diagnosed as having aborted (white bars) or not (black bars). Grey bars show the distribution of *q* among full-term and postpartum *G. pallidipes* sampled in burrows.

Conversely, when *q* < 0.82, no female *G. pallidipes* with an empty uterus was found to have produced a pupa. Accordingly, 0.82 was taken as the desired cut-off for diagnosing an abortion among *G. pallidipes* that had ovulated at least once, but where the uterus was empty. For cases where missing data made it impossible to calculate this quotient, an abortion was diagnosed if *al* < 1.30 mm. For *G. m. morsitans*, where sample sizes were much smaller, it was more difficult to define a value of *q* that provided such a sharp cut-off. As a first approximation, we use the value of 0.82 found in *G. pallidipes*.

When this cut-off was used to diagnose abortions in *G. pallidipes* caught in traps, 84% of flies with empty uteri were diagnosed as not having aborted. The distribution of the values of *q* for these flies forms a tightly distributed unimodal peak around a value of *q* ≈ 1.0 (Fig, 4B). The remaining 16%, which are diagnosed as abortions, take values of *q* that are approximately uniformly distributed on the interval [0.2 – 0.82] (Fig. 4B). Note that the values of *q* for full-term, and postpartum, *G. pallidipes* from burrows show a very similar distribution to those flies from traps where no abortion was diagnosed. Notice that the value of *q* = 0.82 is quite close to the peak associated with flies that are definitely postpartum (Figs. 3C, D, 4B) and might thus be slightly biased towards counting births as abortions. If we set the cut-off *q* at a level lower than 0.82, the estimated abortion rate would be even further reduced.

### Factors affecting the proportion of flies diagnosed as having had a recent abortion Sampling duration

A preliminary analysis showed that, where traps were cleared at intervals of more than 4 hours, *G. pallidipes* exhibited about double the proportion of abortions compared with flies from traps cleared more frequently (Table 1). Accordingly, all further analyses are restricted to sampling periods not exceeding 4 hours.

**Table 1.** Variation in the estimated numbers of abortion in *G. pallidipes* females as a function of sample duration, for flies caught in odour-baited traps. Percentages in parentheses. Pearson chi-squared (df = 4) = 20.6; P < 0.0001

| Sample duration |  |  |  |
| --- | --- | --- | --- |
| (minutes) | No abortion | Abortion | Total |
| 30 | 81,251 (99.31) | 565 (0.69) | 81,816 |
| 60 | 15,429 (99.57) | 66 (0.43) | 15,495 |
| 120 | 11,508 (99.44) | 65 (0.56) | 11,573 |
| 240 | 778 (99.49) | 4 (0.51) | 782 |
| 480 | 2,000 (98.96) | 21 (1.04) | 2,021 |
| Total | 110,966 (99.35) | 721 (0.65) | 111,687 |

### Capture method

Among nearly 110,00 *G. pallidipes* females caught in odour-baited traps, only 0.64% were diagnosed as having suffered a recent abortion (Table 2). The proportions were nearly 8 times higher for flies caught using the VET or other electric nets. This reflects abortions resulting from the capture procedure, as opposed to those that occur naturally: flies were observed to abort in the collection cages or tubes after being killed on an electric net. Samples from electric nets do not thus provide reliable estimates of natural rates of abortion and, accordingly, no further analyses were carried out on these data.

**Table 2.** Variation between sampling methods in the estimated percentage of abortions in *G. pallidipes* and *G. m. morsitans*

|  | *G. pallidipes* |  | *G. m. morsitans* |  |
| --- | --- | --- | --- | --- |
| Method | Abortions/Total | % abort (95% ci) | Abortions/Total | % abort (95% ci) |
| Traps | 700/109,667 | 0.64 (0.59 - 0.69) | 60/5992 | 1.00 (0.76 - 1.29) |
| VET | 431/8768 | 4.92 (3.47 - 4.85) | 170/5045 | 3.37 (2.89 - 3.91) |
| Targets | 138/3347 | 4.12 (3.47 - 4.85) | 10/338 | 2.96 (2.43 - 5.37) |
| Refuge | 216/10,658 | 2.03 (1.77 - 2.31) | 94/4118 | 2.28 (1.85 - 2.79) |
| Burrows | 17/1197 | 1.42 (0.83 - 2.26) | 23/472 | 4.87 (3.11 - 7.22) |
| Ox | 4/767 | 0.52 (0.14 - 1.33) | 5/1214 | 0.41 (0.13 - 0.96) |
| Total | 1,505/134,401 | 1.12 (1.06 - 1.18) | 362/17,179 | 2.11 (1.90 - 2.33) |

The above problem was not observed with flies caught in odour-baited traps, refuges and burrows. Nonetheless, the proportion of abortions among *G. pallidipes* sampled using the latter two methods was 2-4 times as high as that for flies from odour-baited traps (Table 2). It seemed possible that these results could reflect the fact that refuges and burrows were generally only operated in the hot dry months (September – November), when abortion rates are higher than at other times of the year (Hargrove, 1999a). Preliminary analysis indicated, however, that abortion rates for flies from traps were lower than for refuges (and burrows) in every month of the year including the hot months. Accordingly, the results for the various methods are considered separately. Abortion rates among smaller numbers of *G. pallidipes* caught from a bait ox were not significantly different from those obtained from trapped flies.

### Processing delay

Odour-baited traps were generally operated in the afternoon, when tsetse activity is highest at Rekomitjie (Hargrove & Brady, 1992). In the current study 97.4% of flies from traps were caught after 12:00h and, accordingly, a relatively small proportion (3.2%) of the flies could be dissected on the day they were captured; 89.2% of the flies were dissected on the following day, with a final 7.6% being dissected two days after they were collected. The possibility arises that females could abort after they were captured but before they were dissected. There was, however, no significant effect of the delay between capture and dissection on the estimated proportion of flies diagnosed as having suffered a recent abortion (P>0.05, chi-squared). Accordingly, further analyses were carried out on data pooled on dissection delay.

### Seasonal effects

Figure S1.1 in Supplementary File S1 provides details on monthly values of; A, mean ovarian and wing fray categories. B, mean wing length and catch per day per ox fly round. C, mean percentages of females found to have an empty uterus and those diagnosed as having had a recent abortion. D, mean monthly temperature, and total monthly rainfall, at Rekomitjie.

Changes in the flies’ age structure and size, and in the proportions aborting, or simply observed to have an empty uterus, are all highly correlated with seasonal changes in temperature. At the end of the hot dry season, and just prior to the onset of the rains, flies are smallest and exhibit the highest mean wing fray, and ovarian age categories, and also the highest proportions of empty uteri and of diagnosed recent abortions.

### Trypanosome infection

There was no effect of trypanosome infection, measured as any mouthpart infection, on abortion rates (*P* > 0.1 χ^2^ (1df) for both species; *n* = 34,962 and 1401 for *G. pallidipes* and *G. m. morsitans*, respectively) and, similarly, no effect of infections diagnosed as *T. vivax* or *T. congolense*-type infections.

### Logistic regression analysis of abortion rates sampled using odour-baited traps G. pallidipes

#### Bivariable analysis

Among the variables that were considered likely to affect abortion rates, the following had significant effects; wing fray category (*f*) – LR χ^2^ (5df) = 20.2 (P<0.005), capture month (*cm*) – LR χ^2^ (11df) = 118.1 (P<0.0001), capture year (*cy*) – LR χ^2^ (11df) = 176.4 (P<0.0001), temperature during pregnancy (*tbar91*), the mean daily temperature (*tbar9*) during the 9-days prior to fly capture – LR χ^2^ (1df) = 40.1 (P<0.0001) and wing length (*wlm*) – LR χ^2^ (1df) = 22.9 (P<0.0001). When ovarian category was offered, there was no significant effect overall – LR χ^2^ (6df) = 7.9 (P>0.05), but flies in ovarian category 6 had a significantly lower probability of abortion than those in ovarian category 1 (P<0.05). Temperature and wing length were entered as continuous variables, the remainder as categorical variables.

Odds ratios obtained from the above analyses were used to calculate probabilities of diagnosing recent abortions (Figure S1.2 in the Supplementary File S1). Mean probabilities, viewed as a function of ovarian or wing fray category, varied only between about 0.5 and 0.9% (Figure S1.2A, B). Probabilities decreased, from 1.1% to 0.4% with increasing fly size, as measured by wing length (Figure S1.2C) and increased from 0.4% - 1.4% as *tbar91* increased from 20 – 35°C (Figure S1.2D). Variations with capture month reflected, in part, this effect of temperature – being as low 0.3% in the cooler months and as high as 1.1% in the hot season (Figure S1.2E). Changes in the probability with capture year showed the largest range overall (0.2% - 1.5%); this mainly reflected the very high probabilities observed in 1996 (Figure S1.2E).

#### Multivariable, stepwise, analysis

Ovarian category did not have a significant effect on abortion rates in *G. pallidipes* – LR χ^2^ (6df) = 7.89 (P=0.25 >0.05). Moreover, when added to wing fray, there was no significant improvement to the fit – LR χ^2^ (6df) = 8.59 (P=0.20 >0.05). The fit was improved by adding, to wing fray, the month of capture – LR χ^2^ (11df) = 154.5 (P<0.0001) and the year of capture (LR (10 df) = 196.9; P<0.0001, chi-squared). Adding wing length as a variable did not improve the fit – LR χ^2^ (1df) = 0.53 (P=0.47 >0.05). Adding a measure of temperature (*tbar91*) did result in a significantly improved fit – LR χ^2^ (1df) = 15.0 (P<0.0005), but the coefficient of *tbar91* was less than 1.0. That is to say, the model predicts that the probability of diagnosing a recent abortion *decreases* with increasing temperature. This is likely an artefact arising from the high correlation between capture month and temperature.

An alternative model, which included only *f, wlm* and *tbar91* showed significant effects for all three factors, but the *R*^2^ for the fit was markedly smaller than for the modelling including *f, cm* and *cy*. The results for the latter model are shown in Supplementary File S1, Table S1.3A. The coefficients for the wing fray factor indicate that abortion rates for flies in wing fray category 1 are significantly higher than for those in categories 3 – 6 (Fig. 5).

**Figure 5.**
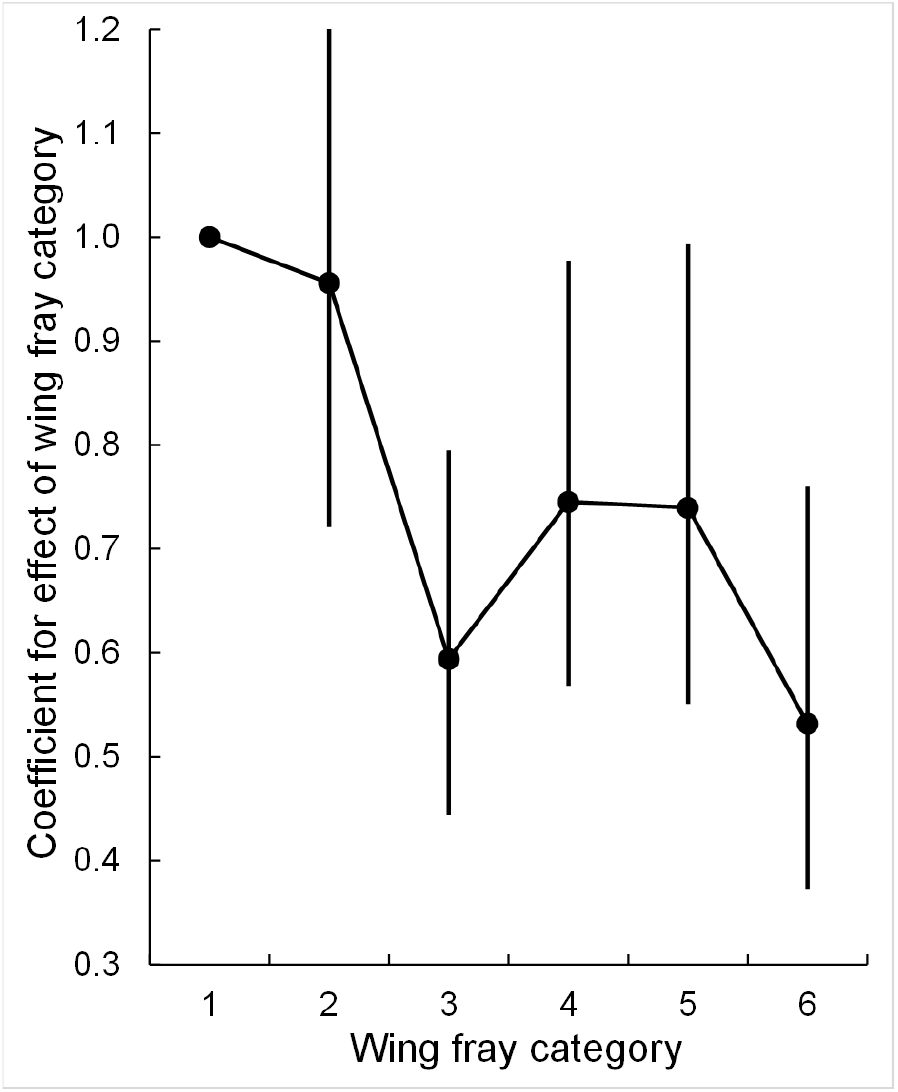
The effect of wing fray category on the probability of diagnosing a recent abortion in female *G. pallidipes*; changes in the coefficient of the wing fray factor included in the logistic regression model detailed in the Supplementary File S1, Table S1.3.

The coefficients for capture month (*cm*) showed a strong peak in December, at the end of the hot-dry season and the onset of the rains (Fig. 6A, C). Changes in mean ovarian and wing fray categories showed closely similar patterns of change (Fig. 6A) so that abortion rates were highest at the time of year when flies were oldest on average, despite the fact that abortion rates were higher in flies in wing fray category 1 (Fig. 5). This effect may be simply attributed to the greater relative effect of increasing temperature compared with increasing age (Supplementary File S1.Tables S1.3A, B).

**Figure 6.**
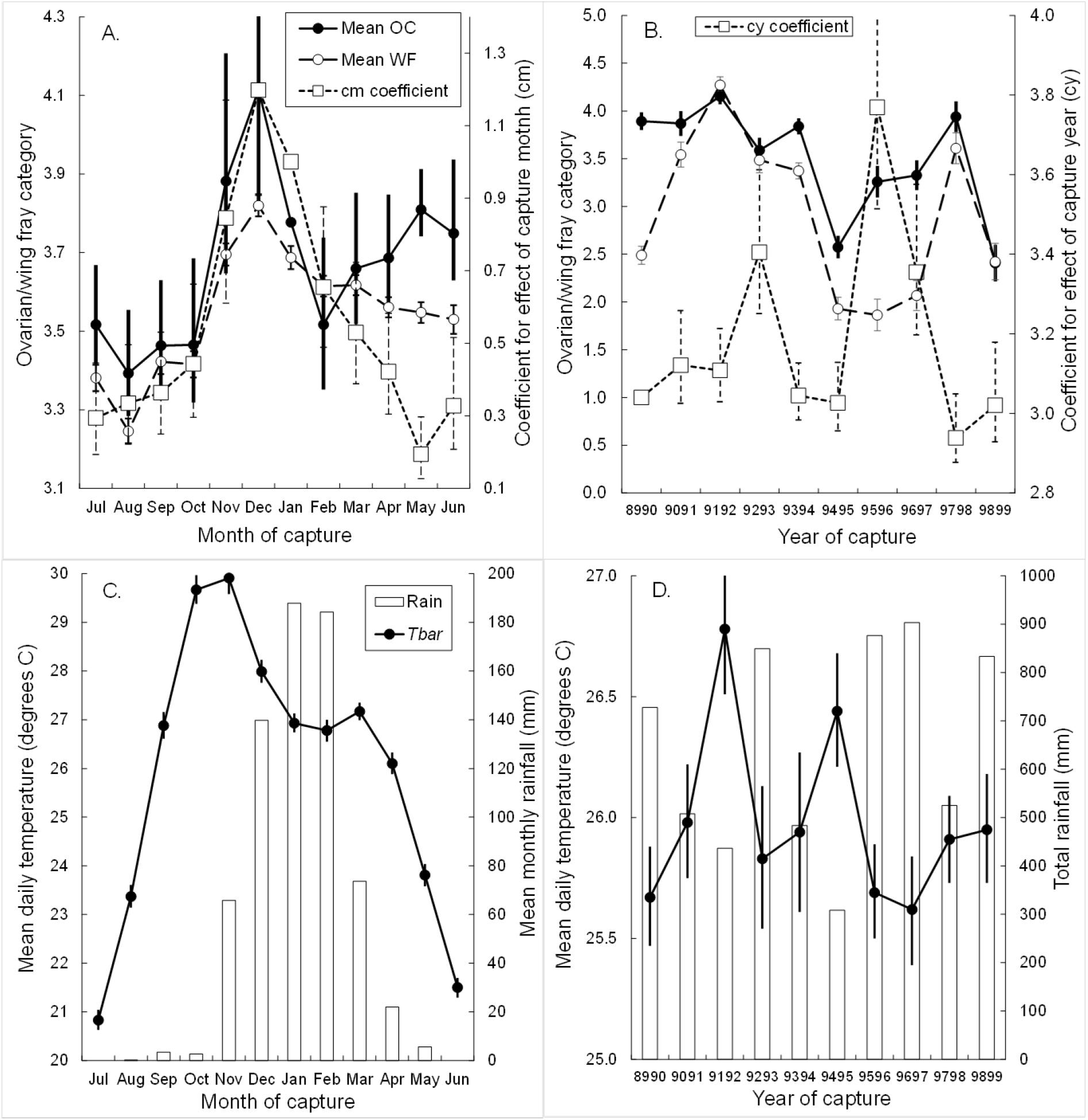
Factors associated with the probability of diagnosing a recent abortion in female *G. pallidipes*. A. Changes, by month of capture, in the mean ovarian (OC) and wing fray (WF) categories, and in the coefficient of the month-of-capture factor included in the logistic regression model detailed in the Supplementary File S1, Table S1.3. January was used as the baseline month for comparison. B. As for A, but by year of capture. The period July 1989 – June 1990 was used as the baseline year for comparison. C. Monthly means (*Tbar*) of daily average temperatures, and mean monthly rainfall, all averaged over the period 1 July 1989 – 30 June 1999. D. As for C, but pooling over years.

For the capture year factor (*cy*), the association between the estimated coefficients, meteorological effects, and measures of age, was less clear. Changes in the coefficients for capture year showed two strong peaks, in 1992/1993 and 1995/1996, which were not obviously related to peaks in mean ovarian and wing fray categories (Fig. 6B). The peaks did, however, occur in the year following particularly adverse weather conditions. Thus, the year 1991/1992 had the highest annual mean temperature recorded at Rekomitjie up to that time – and the second lowest rainfall (Fig. 6B, D). Similarly, the peak in abortion rates in 1995/96 occurred the year after the lowest seasonal rainfall (308) mm over the 50 years when daily rainfall has been recorded at Rekomitjie (Fig. 6B, D).

The peak in the coefficient for *cm* was mirrored by changes in fly sizes (*wlm*), which took minimum values in December and maximum values in May-July during the cool dry season (Fig. 7A). Declines in *wlm* from July-December occur first among flies in ovarian category 0, then in all older flies, consistent with changing wing length being driven via the changes in the sizes of offspring produced. There was no clear pattern for such changes when data were pooled over the year of capture (Fig. 7B).

**Figure 7.**
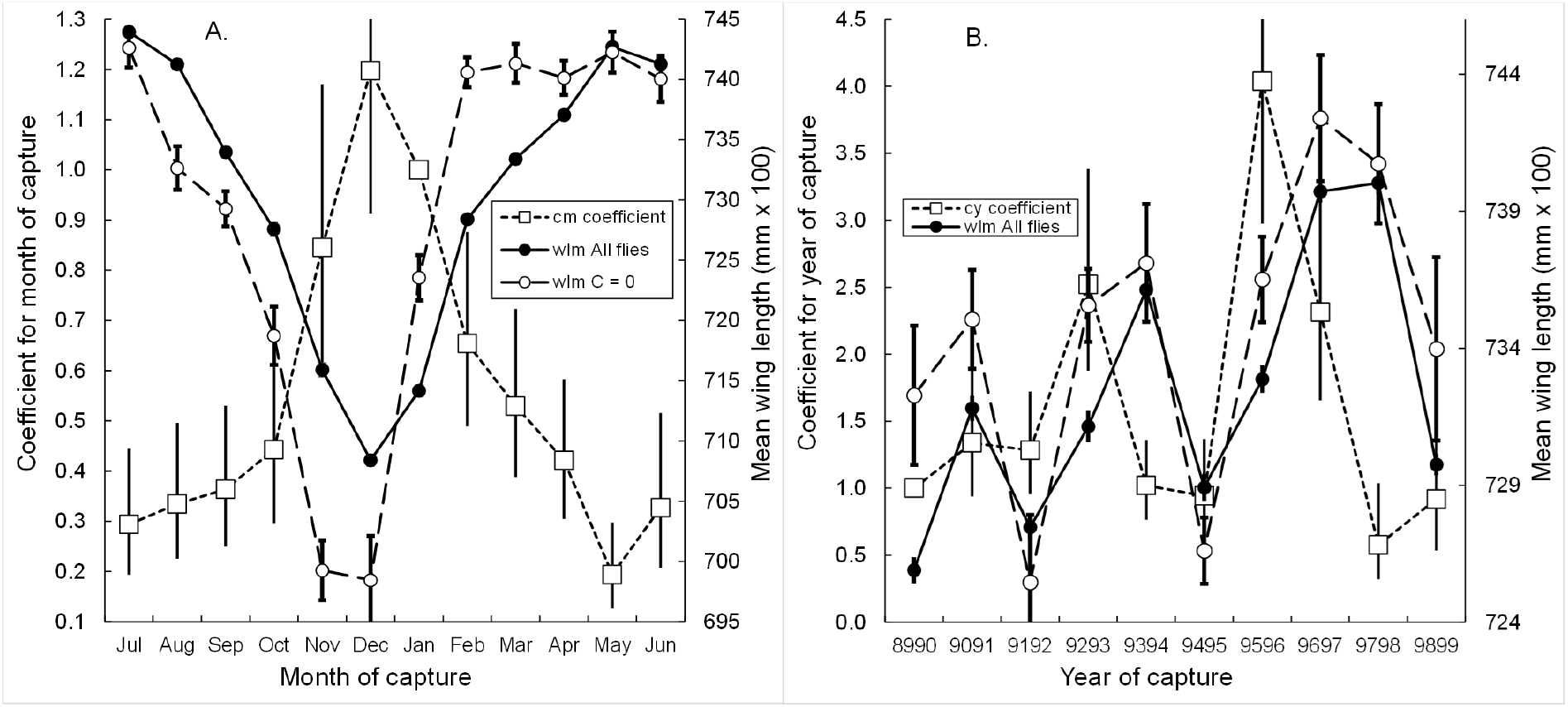
Association between wing length in female *G. pallidipes* and the probability of diagnosing a recent abortion. A. Changes in monthly means of wing lengths among all females dissected, and in the subset young flies, in ovarian category 0, and in the coefficient of capture month (*cm*) in the full logistic regression model. B. As for A. but pooling over year of capture.

#### G. m. morsitans

Analyses of the data for *G. m. morsitans* sampled using odour-baited traps resulted in a qualitatively similar model (Supplementary File S1, Table S1.3B). Given the much smaller numbers of *G. m. morsitans*, patterns were not as clear: nonetheless, as with *G. pallidipes*, abortion rates were highest for flies in wing fray category 1, during the hottest months of the year, and in 1992 and 1996.

#### Predicted probabilities of abortion in G. pallidipes

Probabilities of observing a recent abortion in *G. pallidipes* were calculated from logistic regression results, using Stata’s “*margin*” command, for varying wing fray category and for different capture months. For flies sampled in May the probability was <0.3% and declined only slightly with wing fray category. For flies sampled in December, the probabilities were much higher overall than in May, were 6-times as high for flies in wing fray category 1 and decreased more rapidly with increasing wing fray (Fig. 8A).

**Figure 8.**
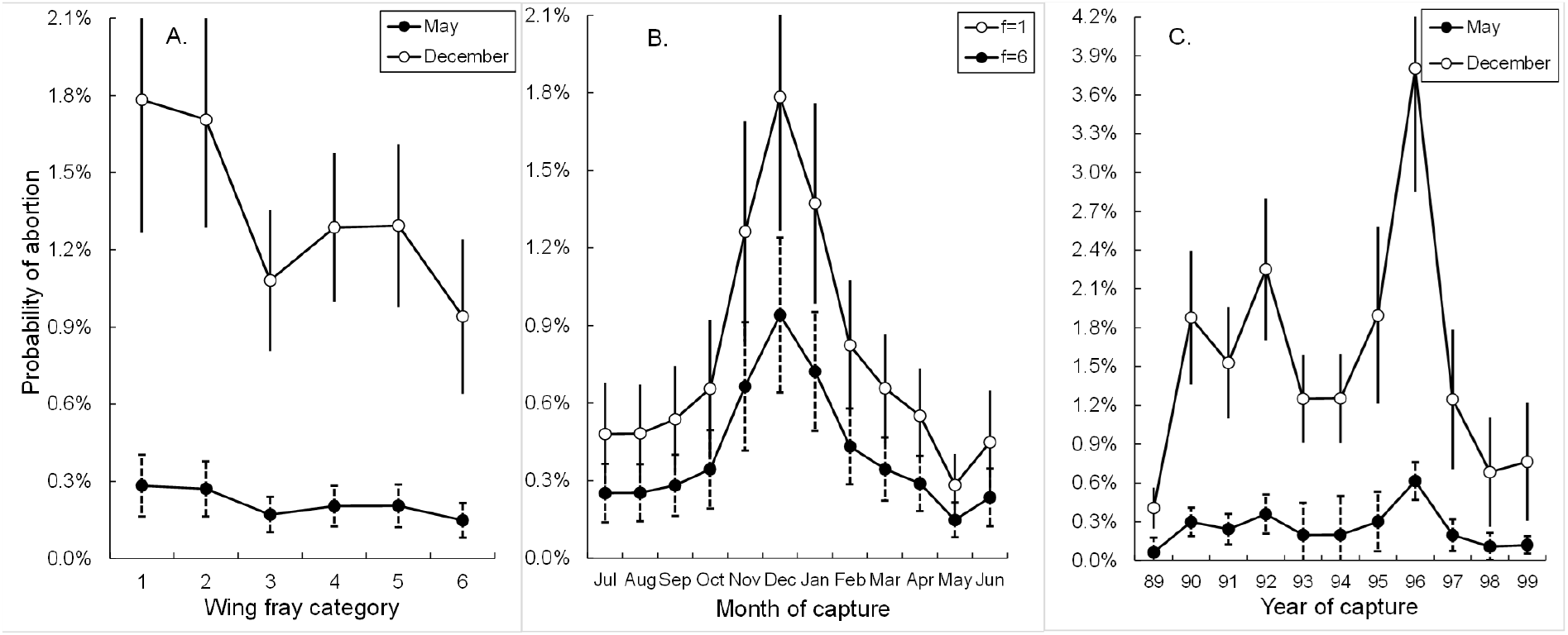
Probabilities of observing a recent abortion in *G. pallidipes* as functions. A. Wing fray category, for flies captured in May or December. B. Month of capture, for flies in wing fray category 1 or 7. C. Year of capture, for flies in wing fray category 1 or 7. In A and B the probabilities are adjusted to the mean of the year of capture, in C to the mean wing fray category.

Similar calculations were carried out to assess changes in abortion probability with capture month, as a function of the wing fray of the flies dissected. There was again a sharp peak in abortions at the end of the hot dry season, with the peak twice as high for flies in wing fray category 1 than for those in category 6 (Fig. 8B). The variation in abortion probability with year of capture depended strongly on the month of the year when the analysis was carried out. For flies captured in May, the probability varied little from a mean level of about 0.3%. For December the overall level was higher, with most years showing probabilities of more than 1%, and was also more variable (Fig. 8C).

#### Abortion rates for flies sampled using artificial refuges

Cross tabulations showed that while only 236/26,075 (0.91%, 0.79% - 1.03%) of *G. pallidipes* caught in odour-baited traps had suffered a recent abortion, the proportion was 2.5 times as high – 212/9382 (2.26%, 1.97% - 2.58%) – when flies were sampled in artificial refuges. For *G. m. morsitans* the corresponding figures were 12/1370 (0.88%, 0.45% - 1.53%) for odour-baited traps and 62/4007 (1.55%, 1.19% - 1.98%) (P>0.05, chi-squared for the difference between methods). For both species the analyses were applied only to flies sampled in September – December when flies were sampled using both traps and artificial refuges.

#### Proportion of female tsetse with empty uteri

Among tsetse of both species, sampled using odour-baited traps, the proportions diagnosed as having suffered an abortion was <1% (see above); the proportions of all flies found on dissection to have an empty uterus, regardless of abortion status, was much higher for all sampling methods (Supplementary File S1, Table S1.5). Thus, among females that had ovulated at least once, the proportions among trapped flies were 4.01% (3.90% - 4.13%, *n* = 109,706) for *G. pallidipes* and 2.52% (2.14% - 2.95%, *n* = 5995) for *G. m. morsitans*. The proportions among flies sampled from refuges were, again, significantly higher; 12.69% (12.07% - 13.34%, *n* = 10,674) for *G. pallidipes* and 14.90% (13.82% - 16.02%, *n* = 4122) for *G. m. morsitans*.

Multiple logistic regressions were carried out, using as the dependent variable the proportion of females that had ovulated at least once, and had an empty uterus, but where *q*>0.82 – such that no abortion was diagnosed. For both *G. pallidipes* and *G. m. morsitans* the above proportion was significantly higher for flies in ovarian category 1 than for all older flies and also decreased with increases in the length of a fly’s wing. As with the proportion of abortions, the proportions of flies with empty uteri were highest in October and November (Supplementary File S1, Tables S1.6A, B).

When the above regression analyses were repeated for flies captured in refuges, flies were only captured between September and December. The same qualitative effects were seen as for flies sampled using traps; the proportion of flies where the uterus was empty was significantly higher in flies in ovarian category 1 than in all older flies and was highest in October and November (Supplementary File S1, Tables S1.7A, B).

## Discussion

### Abortions and premature births

A major aim of this study was to arrive at a satisfactory explanation for the difference between the large proportions of females with empty uteri, and the markedly smaller proportion diagnosed as having suffered an abortion. The concern is that the rule devised here for diagnosing abortions is missing a significant fraction of these events. It was suggested that, among females with empty uteri – captured late in the pregnancy cycle – significant proportions might have aborted earlier in that cycle. Consideration of Figure 4B suggests, however, that this is not a major concern. The frequency of flies where *q* < 0.82 is so uniformly low that they would make little contribution to the relatively huge peak of flies where *q* ≥ 0.82.

A more likely, though only partial, explanation for the discrepancy between the percentages of abortions and of all empty uteri, is simply that the above rule makes too sharp a distinction between a fly that has suffered an abortion, on the one hand, and a fly that has deposited a “normal” larva. The results of the burrows experiment, and other published literature, make it clear that, even where a fly has produced an apparently viable larva, the viability of the resulting adult can be highly variable. Studies on perinatal mortality in tsetse show that there can be very heavy selection pressure against small flies (Jackson, 1948; Bursell & Glasgow, 1960; Phelps & Clarke, 1974; Dransfield et al., 1989). Thus, while a fly that produces a small pupa – from which emerges a living adult – cannot be said to have aborted, the chances of survival of that fly may be much diminished, particularly in extreme weather conditions (Phelps & Clarke, 1974). The dry weights of *G. pallidipes* pupae deposited in artificial warthog burrows varied between about 8 and 18 mg (Hargrove et al., 2018). The smaller pupae in this sample, were not classified as abortions using the rule developed here, but should be classified as premature births. There is thus a grey area between abortions and premature births that is not illuminated by the present classification.

Similar problems occur in all animals, including humans, where a distinction is made between spontaneous abortions and premature births. Howson et al. (2013) highlight the adverse consequences of the latter. Modern medical science can mitigate an impressive proportion of these consequences where, in the past, the probability of survival of preterm babies was much reduced. For tsetse, there is never any such mitigation – except that the female can deposit her larva in a position where it will have climatic conditions that optimise its chances of emerging as a healthy adult.

### Delay between larviposition and ovulation

The preceding section suggests that the relatively large proportion of flies with empty uteri, which have not apparently aborted, may be ascribed in part to a significant level of premature births in tsetse. An additional factor is simply that females in this field study exhibited greater delays between larviposition and subsequent ovulation than reported for laboratory flies. The reasons for the difference cannot be unequivocally resolved with current data – but are consistent with the idea that, where either abortions or premature births have occurred, this is symptomatic of flies that have experienced physiological stress during pregnancy, particularly during inclement climatic conditions. It would not then be surprising if ovulation were delayed while the female rebuilt her nutritional reserves. However, even for flies captured in burrows that were seen to produce a large – presumably full-term and healthy – pupa, the delay in ovulation was much greater than reported by Denlinger & Ma (1974). This could relate to the difference in food availability between field and laboratory tsetse. For the former, feeding is a potentially hazardous and physiologically costly exercise (Randolph et al., 1992; Hargrove & Williams, 1995), whereas laboratory flies feed without danger and with little cost. Postpartum flies in the field, unsurprisingly, have low levels of residual fat (Hargrove & Muzari, 2015). It would not be surprising, therefore, if field flies delayed ovulation until they had secured a further meal.

### Factors affecting proportions of abortions and empty uteri

The close similarity, for both species of tsetse studied here, between the factors associated with an empty uteri – whether or not an abortion is diagnosed – is consistent with these flies having suffered physiological stress in their recent past. It is thus unsurprising that the proportions are highest in the the hot dry season, when adult mortalities are also highest (Hargrove, 2001a, b), and also in years characterised by extreme weather events.

What is more interesting is that, for flies sampled using either odour-baited traps or refuges, the proportions of empty uteri in both species are always highest for flies in the youngest age category, whether age is gauged by ovarian age or wing fray category. Similarly, egg lengths are significantly shorter in these age categories. These results are consistent with laboratory findings (Lord et al. 2021) and with the predictions of an evolutionary model (Barreaux et al., 2022). The field results provide no support, however, for the laboratory finding that abortion rates increase among the oldest females – *cf* Fig. 5, above, with Fig. 2 in Lord et al. (2021). Similarly, the theoretical prediction (Barreaux et al., 2022) that pupal size should decrease in older tsetse is not supported by field results (English et al., 2016; Hargrove et al., 2018). Barreaux et al. (2022) noted the danger of relying solely on results based on laboratory flies and the current work reinforces that concern.

### Differences between samples from refuges and odour-baited traps

The large and statistically significant difference in the frequencies of empty uteri, and of diagnosed abortions, between samples from refuges and odour-baited traps, is not readily explicable. The two systems have quite different sampling biases; stationary odour-baited traps select flies that are at the end of their hunger cycle and the flies are characterised by low fat and haematin levels, whereas tsetse with a broad spectrum of physiological states enter refuges in response to the stresses associated with high temperatures (Hargrove, 1999b, c). If, as suggested above, flies that recently aborted or had undergone premature birth are already physiologically stressed, then these flies might show elevated rates of entering refuges.

A further possible factor is that refuge samples have routinely been taken at 1400h at Rekomitjie, whereas the odour-baited traps were generally operated between about 1600 and 1800h in September - December. Flies that deposit larvae prior to 1400h would be less likely to have ovulated when sampled in refuges than later in the afternoon when captured in a trap. Further work will be required to elucidate this point. It may be concluded, however, that whether the true abortion rate is better reflected by the figures suggested by the trap or the refuge data, that rate is unlikely to be much higher than a modest 2% per pregnancy.

## Acknowledgements

I thank Sinead English for extensive discussions of, and suggestions for, this paper. I am indebted to Dr William Shereni for his continued interest in this work, and for access to the facilities at Rekomitjie Research Station, Zambezi Valley, Zimbabwe. I particularly thank the dissection team at Rekomitjie, and the supervisory role played by Mr Pio Chimanga.

## Availability of data and materials

All data used in this publication are available from the author on reasonable request.

## Ethics approval

Not applicable.

## Consent for publication

The author gives consent for this publication.

