## Supplementary File S1 for "Improved estimates of abortion rates in tsetse (*Glossina* spp)": Supplementary File S1 Abortion rates in tsetse Hargrove 2022 Med Vet Ent.docx

**J. W. Hargrove^*^**

SACEMA, University of Stellenbosch, Stellenbosch, South Africa.

**Supplementary File S1**

**Table S1.1** Details of all tsetse produced in samples of flies collected at Rekomitjie Research Station between October 1988 and December 1999.

|  | *G. m. morsitans* | | *G. pallidipes* | |
| --- | --- | --- | --- | --- |
|  | Numbers | Percent | Numbers | Percent |
| Total | 24,390 | 12.10% | 177,155 | 87.90% |
| Laboratory flies | *13* | 0.05% | *374* | 0.21% |
| *Field flies* |  |  |  |  |
| Pupae | 1134 | 4.65% | 510 | 0.29% |
| Males | 1775 | 7.28% | 15,965 | 9.03% |
| Females | 21,468 | 88.07% | 160,306 | 90.68% |
| *Sampling methods – females* |  |  |  |  |
| Refuges | 5083 | 23.68% | 13,048 | 8.14% |
| Electric nets | 7290 | 33.96% | 16,768 | 10.46% |
| Mechanical traps | 6666 | 31.05% | 126,639 | 79.00% |
| Burrows | 906 | 4.22% | 2643 | 1.65% |
| Bait ox | 1523 | 7.09% | 1208 | 0.75% |
| *Analyses performed – females* |  |  |  |  |
| Nutrition only | 340 | 1.58% | 1915 | 1.19% |
| Infection only | 2 | 0.01% | 60 | 0.04% |
| *Full ovarian dissection* |  |  |  |  |
| *Outcomes* |  |  |  |  |
| Damaged prior to dissection | 1203 | 5.60% | 2975 | 1.86% |
| Damaged during dissection | 219 | 1.02% | 802 | 0.50% |
| Major abnormalities | 86 | 0.40% | 326 | 0.20% |
| Complete | 19,618 | 91.38% | 154,228 | 96.21% |
| *Ovarian category (c)* |  | *Of complete ovarian dissections* |  |  |
| 0 | 1835 | 9.35% | 12,470 | 8.09% |
| 1 | 3316 | 16.90% | 22,925 | 14.86% |
| 2 | 3141 | 16.01% | 22,878 | 14.83% |
| 3 | 2399 | 12.23% | 19,695 | 12.77% |
| 4+4*n* | 3594 | 18.32% | 27,940 | 18.12% |
| 5+4*n* | 2792 | 14.23% | 22,897 | 14.85% |
| 6+4*n* | 1757 | 8.96% | 16,342 | 10.60% |
| 7+4*n* | 784 | 4.00% | 9081 | 5.89% |
| *Uterine content*  *(1≤ c ≤ 7+4n)* |  |  |  |  |
| Empty | 1479 | 8.32% | 7850 | 5.54% |
| Egg | 6882 | 38.70% | 71,079 | 50.14% |
| L1 larva | 3909 | 21.98% | 30,720 | 21.67% |
| L2 larva | 4113 | 23.13% | 29,098 | 20.53% |
| L3 larva | 1390 | 7.82% | 2,939 | 2.07% |
| Undetermined | 10 | 0.06% | 72 | 0.05% |

**Table S1.2A.** Multiple regression analysis of egg length in *G. pallidipes* as a function of maternal wing length (*wlm*), mean maximum temperature during pregnancy (*tmax91*), capture month (*cm*) and year (*cm*), and ovarian category (*c*). Rows of zeroes indicate the reference level for categorical variables.

**.** **regress ulm wlm tmax91 i.cm ib89.cy i.c if g==2 & s==2 & mdpall2<=78 & u==1 & ulm~=. & wlm~=. & ulm>=130 & ulm<=210 & wlm>=615 & wlm<=830 & c>0 & c<8**

```
      Source |       SS           df       MS      Number of obs   =    67,057
-------------+----------------------------------   F(30, 67026)    =    677.97
       Model |   894588.96        30   29819.632   Prob > F        =    0.0000
    Residual |  2948041.81    67,026  43.9835557   R-squared       =    0.2328
-------------+----------------------------------   Adj R-squared   =    0.2325
       Total |  3842630.77    67,056  57.3048014   Root MSE        =     6.632
```

**------------------------------------------------------------------------------**

```
         ulm |      Coef.   Std. Err.      t    P>|t|     [95% Conf. Interval]
-------------+----------------------------------------------------------------
         wlm |   0.024275   0.001118    21.70   0.000     0.022083    0.026467
      tmax91 |  -0.614609   0.015388   -39.94   0.000    -0.644768   -0.584449

          cm |
   January   |          0          0        0       0            0           0
   February  |   0.225256   0.129054     1.75   0.081    -0.027690    0.478201
      March  |   0.924041   0.126205     7.32   0.000     0.676680    1.171402
      April  |   0.955762   0.123939     7.71   0.000     0.712841    1.198682
        May  |   0.335407   0.132216     2.54   0.011     0.076262    0.594550
       June  |  -1.277869   0.155236    -8.23   0.000    -1.582131   -0.973607
       July  |  -2.312210   0.157785   -14.65   0.000    -2.621469   -2.002951
     August  |  -2.825018   0.142529   -19.82   0.000    -3.104375   -2.545661
  September  |  -2.557041   0.130959   -19.53   0.000    -2.813721   -2.300361
    October  |  -4.142197   0.135559   -30.56   0.000    -4.407892   -3.876502
   November  |  -6.060308   0.145668   -41.60   0.000    -6.345816   -5.774799
   December  |  -3.074898   0.135189   -22.75   0.000    -3.339868   -2.809928

          cy |
         88  |   1.572704   0.164216     9.58   0.000     1.250840    1.894567
         89  |          0          0        0       0             0           0
         90  |   0.096574   0.105098     0.92   0.358    -0.109418    0.302567
         91  |  -0.071982   0.106207    -0.68   0.498    -0.280148    0.136185
         92  |  -0.899098   0.102679    -8.76   0.000    -1.100349   -0.697847
         93  |   1.223827   0.095312    12.84   0.000     1.037015    1.410639
         94  |   0.156066   0.097525     1.60   0.110    -0.035084    0.347216
         95  |   0.970499   0.146338     6.63   0.000     0.683676    1.257321
         96  |   1.871042   0.131762    14.20   0.000     1.612789    2.129295
         97  |   3.276960   0.161188    20.33   0.000     2.961032    3.592888
         98  |   1.605363   0.145930    11.00   0.000     1.319341    1.891385
         99  |   2.791850   0.164933    16.93   0.000     2.468581    3.115118

           c |
          1  |          0          0        0       0            0           0
          2  |   2.031855   0.092353    22.00   0.000     1.850844    2.212867
          3  |   3.209243   0.096009    33.43   0.000     3.021065    3.397420
          4  |   2.756654   0.090742    30.38   0.000     2.578800    2.934507
          5  |   3.458299   0.094581    36.56   0.000     3.272920    3.643679
          6  |   3.225303   0.100477    32.10   0.000     3.028367    3.422238
          7  |   3.129911   0.118943    26.31   0.000     2.896783    3.363039
             |
       _cons |   171.26     0.930636    184.03  0.000     169.44      173.09
```

**------------------------------------------------------------------------------**

**C:\Users\jhargrove\OneDrive - Stellenbosch University\dataJWH\TSETSE\Dbdat\OD1to27\Abortions\ EggLengthNew2020Gp1.docx**

**Table S1.2B.** Multiple regression analysis of egg length in *G. m. morsitans* as a function of maternal wing length (*wlm*), mean maximum temperature during pregnancy (*tmax91*), capture month (*cm*) and year (*cm*), and ovarian category (*c*). Rows of zeroes indicate the reference level for categorical variables.

```
. regress ulm wlm tmax91 i.cm ib89.cy i.c if g==1 & s==2 & mdpall2<=78 & u==1 & ulm~=. & wlm~=.  & ulm>=120 & ulm<=190 & wlm>=540 & wlm<=730 & c>0 & c<8

      Source |       SS           df       MS      Number of obs   =     6,244
-------------+----------------------------------   F(30, 6213)     =     66.65
       Model |  99387.1027        30  3312.90342   Prob > F        =    0.0000
    Residual |  308837.237     6,213  49.7082307   R-squared       =    0.2435
-------------+----------------------------------   Adj R-squared   =    0.2398
       Total |   408224.34     6,243  65.3891303   Root MSE        =    7.0504

------------------------------------------------------------------------------
         ulm |      Coef.   Std. Err.      t    P>|t|     [95% Conf. Interval]
-------------+----------------------------------------------------------------
         wlm |   0.031841   0.004198     7.58   0.000      0.023611   0.040071
      tmax91 |  -0.504042   0.056450    -8.93   0.000     -0.614704  -0.393381

         cm  |
    January  |          0          0        0       0            0           0
   February  |   0.381100   0.542292     0.70   0.482     -0.681930   1.444230
      March  |   1.091913   0.526505     2.07   0.038      0.059781   2.124046
      April  |   1.880342   0.545607     3.45   0.001      0.810764   2.949919
        May  |   3.104965   0.541124     5.74   0.000      2.044175   4.165755
       June  |   0.085392   0.527857     0.16   0.871     -0.949390   1.120174
       July  |  -0.553742   0.545589    -1.01   0.310     -1.623285   0.515801
     August  |  -0.574012   0.549919    -1.04   0.297     -1.652043   0.504019
  September  |   0.291186   0.514306     0.57   0.571     -0.717032   1.299404
    October  |   -1.77629   0.512940    -3.46   0.001     -2.781831  -0.770750
   November  |  -5.004312   0.571463    -8.76   0.000     -6.124577  -3.884046
   December  |  -3.412084   0.552138    -6.18   0.000     -4.494466  -2.329702

         cy  |
         88  |   2.067390   0.960155     2.15   0.031      0.185153   3.949626
         89  |          0          0        0       0             0          0
         90  |  -1.754430   0.339721    -5.16   0.000     -2.420400  -1.088460
         91  |  -0.179643   0.305183    -0.59   0.556     -0.777907   0.418622
         92  |  -0.118704   0.354859    -0.33   0.738     -0.814351   0.576943
         93  |  -0.337574   0.352518    -0.96   0.338     -1.028631   0.353483
         94  |  -1.728927   0.380510    -4.54   0.000     -2.474859  -0.982996
         95  |  -1.773713   0.577124    -3.07   0.002     -2.905076  -0.642350
         96  |  -0.971321   0.541094    -1.80   0.073     -2.032052   0.089411
         97  |   1.022825   0.779510     1.31   0.190     -0.505284   2.550934
         98  |  -1.876109   0.543644    -3.45   0.001     -2.941840  -0.810378
         99  |  -1.428811   1.049029    -1.36   0.173     -3.485271   0.627650

          c  |
          1  |          0          0        0       0             0          0
          2  |   2.303815   0.303720     7.59   0.000      1.708420   2.899210
          3  |   2.800552   0.320513     8.74   0.000      2.172236   3.428869
          4  |   2.981288   0.301397     9.89   0.000      2.390446   3.572130
          5  |   2.930712   0.320811     9.14   0.000      2.301812   3.559613
          6  |   2.967034   0.348997     8.50   0.000      2.282879   3.651189
          7  |   3.139217   0.449076     6.99   0.000      2.258873   4.019561
             |
       _cons |   151.6151   3.063297    49.49   0.000      145.61     157.6203
------------------------------------------------------------------------------
```

**C:\Users\jhargrove\OneDrive - Stellenbosch University\dataJWH\TSETSE\Dbdat\OD1to27\Abortions\ EggLengthNew2020Gp1.docx**

**Table S1.3A.** Logistic regression analysis of the probability of observing a recent abortion in *G. pallidipes* as a function of wing fray category (*f*), and capture month (*cm*) and year (*cy*). Flies sampled using odour-baited traps, cleared at intervals not exceeding 240 minutes.

```
. logistic abortnew2 i.f i.cm i.cy if durmin<=240 & mdpall2==44 & g==2 & s==2 & c>0 & c<8 & f>0 & f<7 & cy>88

Logistic regression                             Number of obs     =    106,233
                                                LR chi2(26)       =     372.00
                                                Prob > chi2       =     0.0000
Log likelihood = -3961.9799                     Pseudo R2         =     0.0448

------------------------------------------------------------------------------
   abortnew2 |   Odds Ratio Std. Err.       z   P>|z|     [95% Conf. Interval]
-------------+----------------------------------------------------------------
          f  |
          1  |   1.0
          2  |   0.955738   0.135416    -0.32   0.749     0.723993   1.261664
          3  |   0.601692   0.087890    -3.48   0.001     0.451895   0.801144
          4  |   0.717825   0.097928    -2.43   0.015     0.549408   0.937869
          5  |   0.721796   0.106798    -2.20   0.028     0.540095   0.964626
          6  |   0.522950   0.093948    -3.61   0.000     0.367749   0.743677
             |
         Cm  |
    January  |   1.0
   February  |   0.596789   0.087556    -3.52   0.000     0.447651   0.795613
      March  |   0.475393   0.074128    -4.77   0.000     0.350208   0.645327
      April  |   0.398035   0.065273    -5.62   0.000     0.288625   0.548917
        May  |   0.204186   0.043177    -7.51   0.000     0.134906   0.309044
       June  |   0.324147   0.072494    -5.04   0.000     0.209109   0.502470
       July  |   0.346983   0.072587    -5.06   0.000     0.230273   0.522847
     August  |   0.348560   0.069875    -5.26   0.000     0.235310   0.516317
  September  |   0.388436   0.074875    -4.91   0.000     0.266220   0.566758
    October  |   0.474606   0.097076    -3.64   0.000     0.317854   0.708661
   November  |   0.919569   0.153816    -0.50   0.616     0.662528   1.276334
   December  |   1.305035   0.182130     1.91   0.056     0.992725   1.715598
             |
         cy  |
         89  |   1.0
         90  |   4.691488   1.008874     7.19   0.000     3.077976   7.150823
         91  |   3.807205   0.840081     6.06   0.000     2.470495   5.867169
         92  |   5.650181   1.153222     8.48   0.000     3.787295   8.429380
         93  |   3.104779   0.672006     5.23   0.000     2.031396   4.745334
         94  |   3.112548   0.658349     5.37   0.000     2.056244   4.711482
         95  |   4.736863   1.157368     6.37   0.000     2.934370   7.646572
         96  |   9.696277   2.034518    10.83   0.000     6.426905  14.628780
         97  |   3.094453   0.851393     4.11   0.000     1.804630   5.306149
         98  |   1.686698   0.600488     1.47   0.142     0.839456   3.389042
         99  |   1.887998   0.648075     1.85   0.064     0.963424   3.699861
             |
       _cons |   0.004318   0.000997   -23.57   0.000     0.002745   0.006790
------------------------------------------------------------------------------
```

**Table S1.3B.** Logistic regression analysis of the probability of observing a recent abort in *G. m. morsitans* as a function of wing fray category (*f*), and capture month (*cm*) and year (*cy*). Flies sampled using odour-baited traps, cleared at intervals not exceeding 240 minutes.

```
. logistic abortnew2 i.f i.cm i.cy if durmin<=240 & mdpall2==44 & g==1 & s==2 & c>0
                     & c<8 & f>0 & f<7 & cy>88 & cy<97

note: 6.f  != 0 predicts failure perfectly		6.f  dropped and 234 obs not used
note: 8.cm != 0 predicts failure perfectly		8.cm dropped and 335 obs not used


Logistic regression                             Number of obs     =      4,824
                                                LR chi2(21)       =      63.33
                                                Prob > chi2       =     0.0000
Log likelihood =  -228.1386                     Pseudo R2         =     0.1219

------------------------------------------------------------------------------
   abortnew2 | Odds Ratio   Std. Err.      z    P>|z|     [95% Conf. Interval]
-------------+----------------------------------------------------------------
          f  |
          1  |   1.0
          2  |   0.656662   0.249156    -1.11   0.268     0.312156   1.381377
          3  |   0.158023   0.099648    -2.93   0.003     0.045915   0.543854
          4  |   0.361488   0.159359    -2.31   0.021     0.152352   0.857708
          5  |   0.355003   0.202755    -1.81   0.070     0.115899   1.087386
          6  |              (empty)
             |
         Cm  |
    January  |   1.0
   February  |   0.086954   0.089870    -2.36   0.018     0.011469    0.659233
      March  |   0.743409   0.399947    -0.55   0.582     0.258994    2.133860
      April  |   0.106645   0.113753    -2.10   0.036     0.013183    0.862721
        May  |   0.419713   0.233628    -1.56   0.119     0.140974    1.249578
       June  |   0.135726   0.145301    -1.87   0.062     0.016650    1.106416
       July  |   0.149339   0.161919    -1.75   0.079     0.017835    1.250477
     August  |              (empty)
  September  |   0.128733   0.104992    -2.51   0.012     0.026030    0.636662
    October  |   0.396140   0.326122    -1.12   0.261     0.078904    1.988837
   November  |   0.624031   0.396701    -0.74   0.458     0.179511    2.169306
   December  |   0.396048   0.280271    -1.31   0.191     0.098942    1.585317
             |
         cy  |
         90  |   1.041742   0.603673     0.07   0.944     0.334582    3.243533
         91  |   0.585671   0.638578    -0.49   0.624     0.069113    4.963049
         92  |   6.538173   4.089479     3.00   0.003     1.918885   22.277360
         93  |   2.744680   1.785196     1.55   0.121     0.767101    9.820436
         94  |   1.186437   0.708585     0.29   0.775     0.368023    3.824850
         95  |   1.746499   1.483595     0.66   0.512     0.330446    9.230742
         96  |   5.125624   3.048026     2.75   0.006     1.597966   16.440910
             |
       _cons |   0.032422   0.019668    -5.65   0.000     0.009874    0.106464
------------------------------------------------------------------------------
```

**Table S1.4A.** Logistic regression analysis of the probability of observing a recent abortion in *G. pallidipes*, sampled using artificial refuges, as a function of ovarian category (*c*), and capture month (*cm*) and year (*cy*).

```
. logistic abortnew2 i.cy i.cm i.c if md==1 & g==2 & s==2 & c>0 & c<8 & f>0 & f<7 & cm>8
Logistic regression                             Number of obs     =      9,027
                                                LR chi2(18)       =      92.30
                                                Prob > chi2       =     0.0000
Log likelihood = -801.45742                     Pseudo R2         =     0.0544
------------------------------------------------------------------------------
   abortnew2 | Odds Ratio   Std. Err.      z    P>|z|     [95% Conf. Interval]
-------------+----------------------------------------------------------------
         Cy  |
         88  |    1.0
         89  |    3.04463   1.744470     1.94   0.052     0.990431    9.35932
         90  |   12.70102   8.788336     3.67   0.000     3.272342   49.29677
         91  |   13.48757   7.763099     4.52   0.000     4.365198   41.67383
         92  |   44.92400  27.011400     6.33   0.000     13.82538  145.97530
         93  |   15.66826   8.920308     4.83   0.000     5.133481   47.82222
         94  |   11.12330   6.647619     4.03   0.000     3.447705   35.88701
         95  |   16.99932  11.237140     4.29   0.000     4.653264   62.10196
         96  |    7.21618   4.303909     3.31   0.001     2.241968   23.22657
         98  |   13.53032   7.766191     4.54   0.000     4.392701   41.67584

         cm  |
  September  |    1.0
    October  |    0.82170   0.184290    -0.88   0.381     0.529428    1.275323
   November  |    1.48693   0.387236     1.52   0.128     0.892512    2.477227
   December  |    2.84777   1.344501     2.22   0.027     1.128836    7.184186

          C  |
          1  |    1.0
          2  |    1.67392   0.503314     1.71   0.087     0.928529    3.017686
          3  |    2.07200   0.616534     2.45   0.014     1.156403    3.712520
          4  |    1.19712   0.370962     0.58   0.562     0.652180    2.197389
          5  |    1.57272   0.487230     1.46   0.144     0.856934    2.886385
          6  |    2.00704   0.658301     2.12   0.034     1.055272    3.817214
          7  |    2.69158   0.984133     2.71   0.007     1.314556    5.511065
       _cons |    0.00118   0.000732   -10.86   0.000     .0003491    0.003980
------------------------------------------------------------------------------
```

**Table S1.4B.** Logistic regression analysis of the probability of observing a recent abortion in *G. m. morsitans*, sampled using artificial refuges, as a function of capture year (*cy*) and the mean temperature (*tbar91*) over the 9-day period prior to capture.

```
. logistic abortnew2 tbar91 i.cy if durmin<=240 & cy>88 & (mdpall2==1|mdpall2==44) & g==1 & s==2 & wlm>=525 & wlm<=725 & c>0 & c<8 & f>0 & f<7 & cm>8 & cy~=97 & cy~=99

Logistic regression                             Number of obs     =      5,377
                                                LR chi2(9)        =      40.83
                                                Prob > chi2       =     0.0000
Log likelihood = -370.22648                     Pseudo R2         =     0.0523
------------------------------------------------------------------------------
   abortnew2 | Odds Ratio   Std. Err.      z    P>|z|     [95% Conf. Interval]
-------------+----------------------------------------------------------------
      tbar91 |   1.13816    0.065902     2.24   0.025     1.016054    1.274941
             |
         cy  |
         98  |   1.0
         90  |   1.988212   1.541255     0.89   0.375     .4351252    9.084709
         91  |   1.148377   .9969792     0.16   0.873     .2094607    6.296028
         92  |   5.073165   2.958388     2.78   0.005     1.617728    15.90935
         93  |   2.244161   1.380292     1.31   0.189      .672225    7.491927
         94  |   5.795148   3.275647     3.11   0.002     1.913955    17.54677
         95  |   8.634716   4.946396     3.76   0.000     2.809554    26.53742
         96  |   2.944111    1.88808     1.68   0.092     .8376689    10.34751
         98  |   4.182049   2.719426     2.20   0.028     1.169191    14.95867
             |
       _cons |   .0000875   .0001561    -5.24   0.000     2.66e-06    .0028851
```

**Table S1.5.** Variation between sampling methods in the percentage of *G. pallidipes* and *G. m. morsitans* where the uterus was empty in females that had ovulated at least once

|  | *G. pallidipes* | | *G. m. morsitans* | |
| --- | --- | --- | --- | --- |
| Method | Empty/Total | % empty (95% ci) | Empty/Total | % empty (95% ci) |
| Traps | 402/109,706 | 4.01 (3.90 - 4.13) | 151/5995 | 2.52 (2.14 – 2.95) |
| VET | 796/8785 | 9.06 (8.47 - 9.68) | 365/5050 | 7.23 (6.53 - 7.98) |
| Targets | 267/3351 | 7.97 (7.07 - 8.93) | 16/338 | 4.73 (2.73 – 7.57) |
| Refuge | 1355/10,674 | 12.69 (12.07 - 13.34) | 614/4122 | 14.90 (13.82 - 16.02) |
| Burrows | 469/1197 | 39.18 (36.40 - 42.01) | 197/472 | 41.74 (37.25 - 46.33) |
| Ox | 10/767 | 1.30 (0.63 - 2.38) | 16/1214 | 1.32 (0.76 – 2.13) |
| Total | 7299/134,480 | 5.43 (5.31 – 5.55) | 1359/17,191 | 7.91 (7.51 - 8.32) |

**Table S1.6A.** Logistic regression analysis of the probability of observing an empty uterus in *G. pallidipes*, sampled using odour-baited traps, as a function of the mean temperature (*tbar91*) over the 9-day period prior to capture, wing length (*wlm*), ovarian category (*c*), and the month (*cm*) and year (*cy*) of capture. The analysis was limited to flies that were diagnosed as not having suffered a recent abortion.

```
. logistic umt wlm i.c i.cm i.cy if abortnew2==0 & durmin<=240 & mdpall2==44 & g==2 & s==2 & c>0 & c<8 & wlm~=.

Logistic regression                             Number of obs     =    109,125
                                                LR chi2(29)       =    2868.74
                                                Prob > chi2       =     0.0000
Log likelihood = -14609.531                     Pseudo R2         =     0.0894

------------------------------------------------------------------------------
         umt | Odds Ratio   Std. Err.      z    P>|z|     [95% Conf. Interval]
-------------+----------------------------------------------------------------
         wlm |   0.9972992   .0007452   -3.62   0.000     0.995840    0.998761
             |
           c |
          1  |   1.0
          2  |   0.629431   0.033204    -8.78   0.000     0.567604    0.697993
          3  |   0.424759   0.026298   -13.83   0.000     0.376220    0.479559
          4  |   0.485805   0.025643   -13.68   0.000     0.438057    0.538757
          5  |   0.462593   0.026150   -13.64   0.000     0.414077    0.516793
          6  |   0.352934   0.024575   -14.96   0.000     0.307911    0.404541
          7  |   0.326349   0.029728   -12.29   0.000     0.272989    0.390139
             |
          cm |
   February  |   1.078117   0.087061     0.93   0.352     0.920300    1.262998
      March  |   1.077760   0.089224     0.90   0.366     0.916335    1.267623
      April  |   0.732727   0.065322    -3.49   0.000     0.615260    0.872621
        May  |   0.647830   0.060654    -4.64   0.000     0.539221    0.778315
       June  |   0.604399   0.068952    -4.41   0.000     0.483299    0.755843
       July  |   0.575539   0.062414    -5.09   0.000     0.465337    0.711841
     August  |   0.430693   0.050188    -7.23   0.000     0.342751    0.541200
  September  |   1.137086   0.100509     1.45   0.146     0.956212    1.352175
    October  |   2.500954   0.203110    11.29   0.000     2.132933    2.932474
   November  |   4.768976   0.361792    20.59   0.000     4.110076    5.533507
   December  |   1.908399   0.156895     7.86   0.000     1.624387    2.242068
             |
          cy |
         89  |    3.73801    1.10316     4.47   0.000     2.096202     6.66571
         90  |   12.61884    3.65400     8.76   0.000     7.153842    22.25869
         91  |   32.27365    9.13538    12.27   0.000     18.53135    56.20684
         92  |   39.84178   11.25945    13.04   0.000     22.89738    69.32530
         93  |   20.80623    5.88709    10.73   0.000     11.94943    36.22761
         94  |   29.21896    8.26200    11.94   0.000     16.78717    50.85715
         95  |   36.12480   10.40967    12.45   0.000     20.53641    63.54573
         96  |   37.73350   10.79592    12.69   0.000     21.53724    66.10953
         97  |   21.99910    6.41298    10.60   0.000     12.42426    38.95286
         98  |   28.50272    8.49074    11.25   0.000     15.89717    51.10375
         99  |   14.86725    4.42351     9.07   0.000     8.297925    26.63739
             |
       _cons |    0.01640    0.00988    -6.82   0.000     0.005033    0.053417
------------------------------------------------------------------------------
```

**Table S1.6B.** Logistic regression analysis of the probability of observing an empty uterus, among *G. m. morsitans* sampled using odour-baited traps, as a function of ovarian category (*c*), and the month (*cm*) and year (*cy*) of capture. The analysis was limited to flies that were diagnosed as not having suffered a recent abortion.

```
. logistic umt wlm i.c i.cm i.cy if abortnew2==0 & durmin<=240 & mdpall2==44 & g==1 & s==2 & c>0 & c<8 & f>0 & f<7 & wlm~=.

Logistic regression                             Number of obs     =      5,940
                                                LR chi2(29)       =      88.39
                                                Prob > chi2       =     0.0000
Log likelihood = -455.23207                     Pseudo R2         =     0.0885

------------------------------------------------------------------------------
         umt | Odds Ratio   Std. Err.      z    P>|z|     [95% Conf. Interval]
-------------+----------------------------------------------------------------
         wlm |   0.995347   0.004793    -0.97   0.3330      .985997    1.004786
             |
           c |
          1  |   1.0
          2  |   0.457072   0.1449271    -2.47   0.014     .2455209    0.850905
          3  |   0.362458   0.1382594    -2.66   0.008     .1716200    0.765505
          4  |   0.420505   0.1305966    -2.79   0.005     .2287774    0.772911
          5  |   0.484939   0.1618535    -2.17   0.030     .2521101    0.932792
          6  |   0.413862   0.1750643    -2.09   0.037     .1806325    0.948233
          7  |   0.386934   0.2367652    -1.55   0.121     .1166238    1.283772
             |
          cm |
    January  |   1.0
   February  |   0.427981   0.2260627    -1.61   0.108     .1519891    1.205134
      March  |   0.556204   0.3003521    -1.09   0.277     .1930112    1.602824
      April  |   0.536662   0.2654898    -1.26   0.208     .2035195    1.415126
        May  |   0.215191   0.1353812    -2.44   0.015     .0627066    0.738471
       June  |   0.228986   0.1829205    -1.85   0.065     .0478459    1.095905
       July  |   0.453945   0.2878471    -1.25   0.213     .1309952    1.573081
     August  |   0.208582   0.1457512    -2.24   0.025     .0530253    0.820488
  September  |   0.674451   0.3109463    -0.85   0.393     .2732221    1.664888
    October  |   1.749386   0.8204394     1.19   0.233     .6977235    4.386197
   November  |   1.642989   0.7729893     1.06   0.291     .6533765    4.131482
   December  |   0.637916   0.3255622    -0.88   0.378     .2346118    1.734509
             |
          cy |
         89  |   1.0
         89  |   0.411425   0.2762906    -1.32   0.186     .1103243    1.534303
         90  |   0.273352   0.2042068    -1.74   0.083     .0632170    1.181980
         91  |   1.127053   0.7738656     0.17   0.862     .2934173    4.329151
         92  |   1.121924   0.8968321     0.14   0.886     .2341741    5.375120
         93  |   0.748638   0.5246544    -0.41   0.680     .1895567    2.956679
         94  |   0.843693   0.5548400    -0.26   0.796     .2324915    3.061697
         95  |   0.712965   0.6186102    -0.39   0.697     .1301713    3.904997
         96  |   1.967113   1.3551030     0.98   0.326     .5098637    7.589346
         97  |   0.979281   0.6991614    -0.03   0.977     .2416512    3.968496
         98  |   0.477369   0.5715716    -0.62   0.537     .0456750    4.989184
         99  |   0.241685   0.2886211    -1.19   0.234     .0232670    2.510491
             |
       _cons |   1.492911   4.617908     0.13   0.8970      .003476    641.1954
------------------------------------------------------------------------------
```

**Table S1.7A.** Logistic regression analysis of the probability of observing an empty uterus in *G. pallidipes*, sampled using artificial refuges, as a function of the fly’s ovarian category (*c*), and the month (*cm*) and year (*cy*) of capture. The analysis was limited to flies that were diagnosed as not having suffered a recent abortion.

```
. logistic umt i.c i.cm i.cy if abortnew2==0 & md==1 & g==2 & s==2 & c>0 & c<8 & cm>8

Logistic regression                             Number of obs     =      8,881
                                                LR chi2(18)       =     468.61
                                                Prob > chi2       =     0.0000
Log likelihood = -2650.3409                     Pseudo R2         =     0.0812

------------------------------------------------------------------------------
         umt | Odds Ratio   Std. Err.      z    P>|z|     [95% Conf. Interval]
-------------+----------------------------------------------------------------
          C  |
          1  |   1.0
          2  |   0.777694   0.085895    -2.28   0.023     0.626319    0.96566
          3  |   0.544493   0.068570    -4.83   0.000     0.425399    0.69693
          4  |   0.505785   0.059527    -5.79   0.000     0.401593    0.63701
          5  |   0.531685   0.066678    -5.04   0.000     0.415820    0.67983
          6  |   0.444569   0.067838    -5.31   0.000     0.329650    0.59955
          7  |   0.505949   0.097749    -3.53   0.000     0.346461    0.73885
             |
          cm |
  September  |   1.0
    October  |   2.213420   0.289086     6.08   0.000     1.713530    2.85914
   November  |   3.647420   0.505434     9.34   0.000     2.779919    4.78563
   December  |   1.488914   0.470699     1.26   0.208     0.801262    2.76672
             |
          cy |
         88  |   1.0
         89  |   6.727036   1.701096     7.54   0.000     4.098043   11.04259
         90  |   9.928819   3.073970     7.41   0.000     5.412089   18.21505
         91  |  11.791200   3.141496     9.26   0.000     6.994795   19.87656
         92  |  36.222050  10.412730    12.49   0.000    20.619520   63.63081
         93  |   7.278564   1.941882     7.44   0.000     4.314690   12.27804
         94  |  21.216540   5.735678    11.30   0.000    12.489950   36.04029
         95  |  15.965080   4.809095     9.20   0.000     8.846412   28.81210
         96  |   7.340251   1.996139     7.33   0.000     4.307562   12.50807
         98  |  10.780050   2.858992     8.97   0.000     6.410215   18.12880
             |
       _cons |   0.008811   0.002485   -16.78   0.000     0.005069    0.015314
------------------------------------------------------------------------------
```

**Table S1.7B.** Logistic regression analysis of the probability of observing an empty uterus in *G. m. morsitans*, sampled using artificial refuges, as a function of the fly’s ovarian category (*c*), and the month (*cm*) and year (*cy*) of capture. The analysis was limited to flies that were diagnosed as not having suffered a recent abortion.

```
. logistic umt i.c i.cm i.cy if abortnew2==0 & md==1 & g==1 & s==2 & c>0 & c<8 & cm>8

Logistic regression                             Number of obs     =      3,465
                                                LR chi2(18)       =     141.26
                                                Prob > chi2       =     0.0000
Log likelihood = -1284.3586                     Pseudo R2         =     0.0521

------------------------------------------------------------------------------
         umt | Odds Ratio   Std. Err.      z    P>|z|     [95% Conf. Interval]
-------------+----------------------------------------------------------------
           c |
          1  |   1.0
          2  |   0.558530   0.091838    -3.54   0.000     0.4046558    0.77092
          3  |   0.534632   0.098341    -3.40   0.001     0.3728076    0.76670
          4  |   0.566804   0.089186    -3.61   0.000     0.4163874    0.77156
          5  |   0.517168   0.088098    -3.87   0.000     0.3703672    0.72216
          6  |   0.417052   0.087650    -4.16   0.000     0.2762458    0.62963
          7  |   0.322192   0.100777    -3.62   0.000     0.1745301    0.59478
             |
          cm |
  September  |   1.0
    October  |   2.331404   0.480189     4.11   0.000     1.557038    3.490887
   November  |   3.700306   0.804160     6.02   0.000     2.416864    5.665300
   December  |   1.674782   0.749158     1.15   0.249     0.6969499   4.024527
             |
          cy |
         88  |   1.0
         89  |   1.155480   0.438853     0.38   0.704     0.5488767   2.432484
         90  |   1.573133   0.723504     0.99   0.325     0.6386848   3.874750
         91  |   1.242952   0.504429     0.54   0.592     0.561061    2.753585
         92  |   5.631583   2.179171     4.47   0.000     2.637871    12.02285
         93  |   2.099313   0.799053     1.95   0.051     0.995607    4.426562
         94  |   4.438119   1.734959     3.81   0.000     2.062744    9.548881
         95  |   2.344114   0.892650     2.24   0.025     1.111315    4.944479
         96  |   1.740722   0.667775     1.44   0.148     0.820714    3.692045
         98  |   1.918733   0.771279     1.62   0.105     0.872679    4.218662
             |
       _cons |   0.049707   0.020417    -7.31   0.000     0.022222    0.111184
------------------------------------------------------------------------------
```

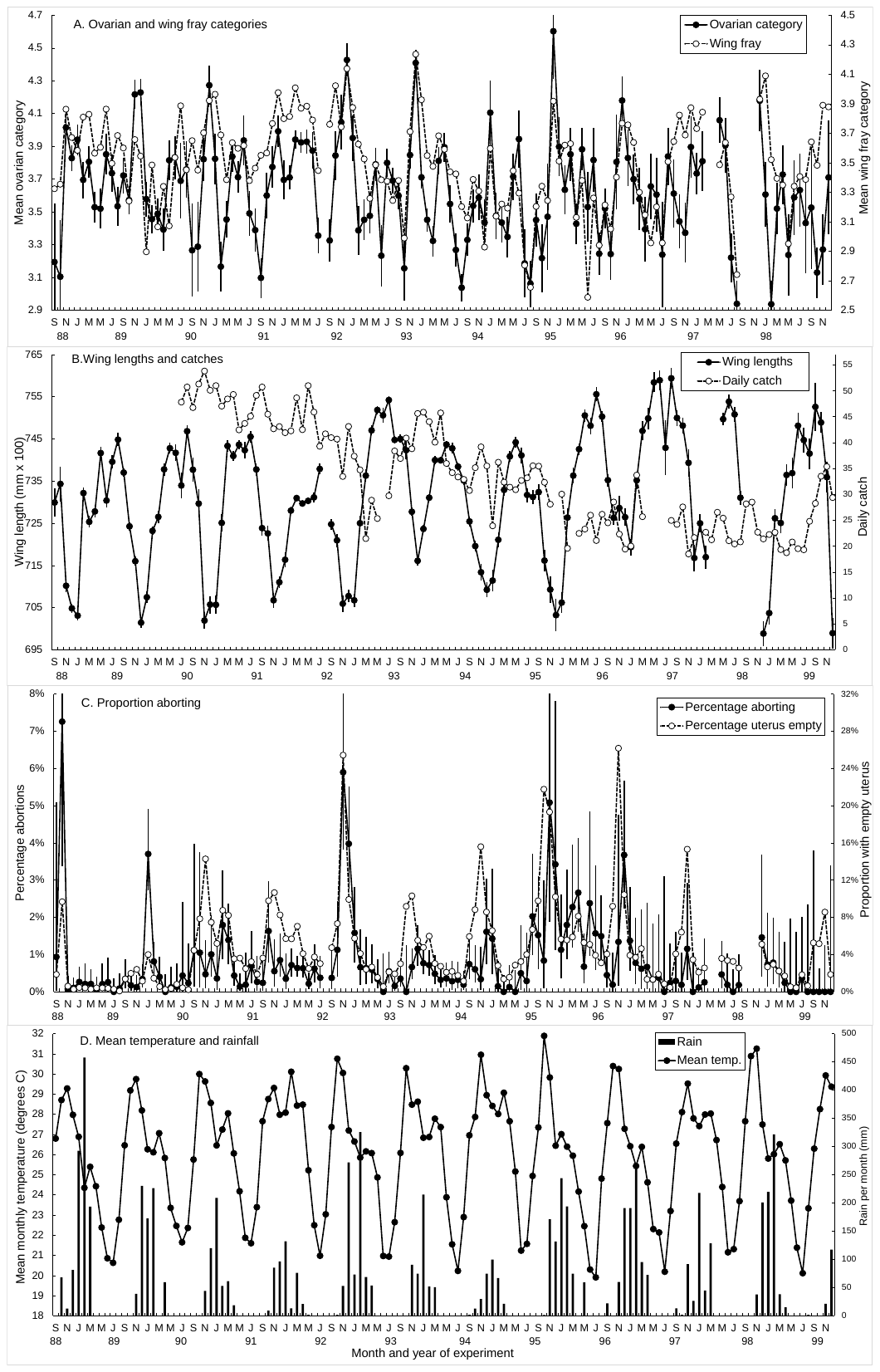

**Figure S1.1.** Overview of monthly changes during September 1988-December 1999 in; A. Mean ovarian, and wing fray, categories. B. Wing lengths and numbers of flies captured per day per ox fly round. C. Percentage of female *G. pallidipes* with empty uterus or diagnosed to have aborted recently. B. Mean temperature and total rainfall per month.

**C:\Users\jhargrove\OneDrive - Stellenbosch University\dataJWH\TSETSE\Dbdat\OD1to27\Abortions\Excel\Logistic regression Best model.xlsx Worksheet: OC-F-WL-Ab by ExpMon**

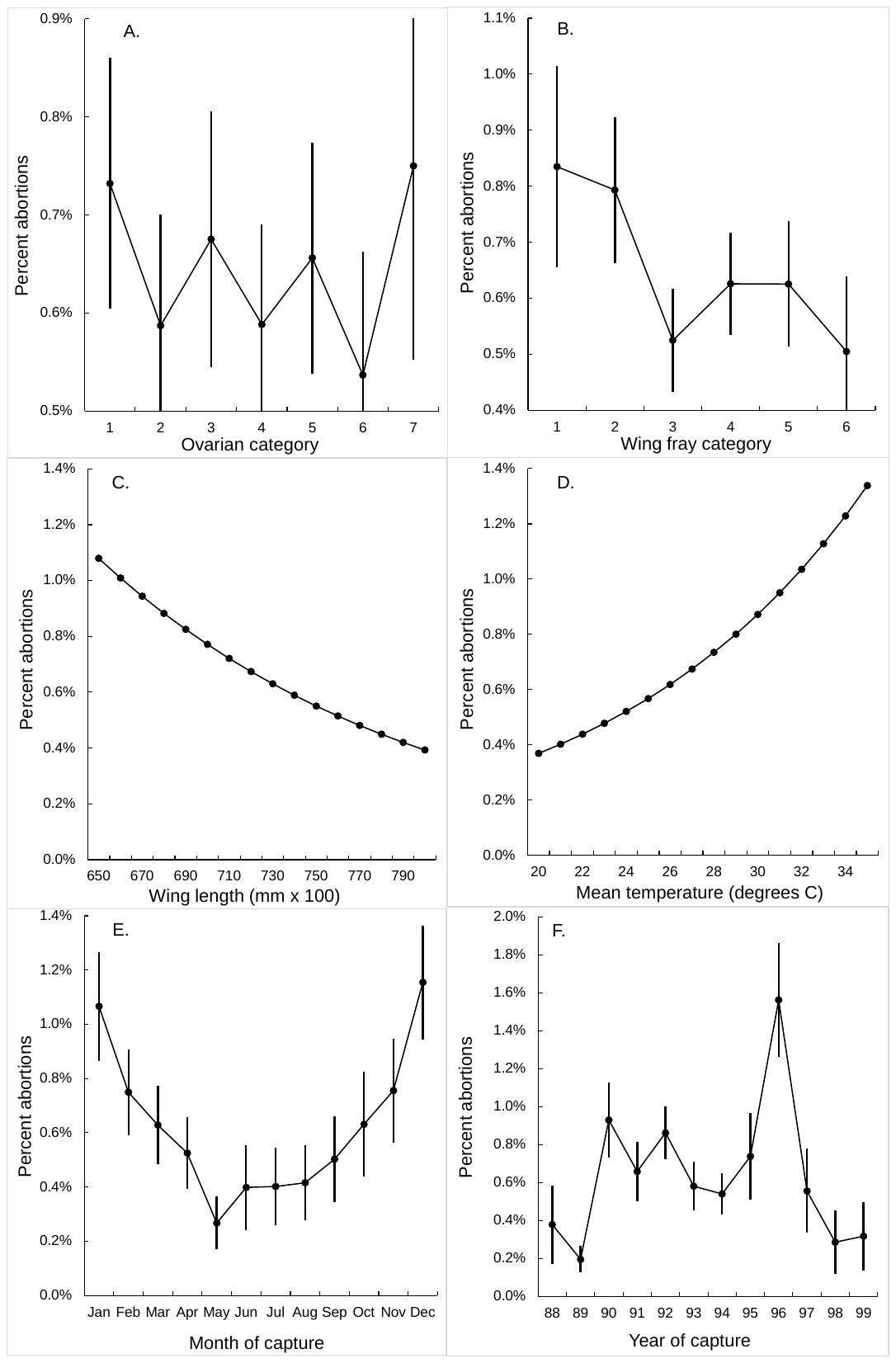

**Figure S1.2.** The percentage of *G. pallidipes* females diagnosed to have experienced a recent abortion, as a function of, A. Ovarian category. B. Wing fray category C. Wing length of the flies sampled and dissected. D. Mean temperature over the nine days prior to the fly being sampled. E. Month of capture. F. Year of capture.

**C:\Users\jhargrove\OneDrive - Stellenbosch University\dataJWH\TSETSE\Dbdat\OD1to27\Abortions\Excel\** **Logistic Regression P Abort vs wf cm cy.xlsx Worksheet Bivariable regression**
